# Deviations from dynamic equilibrium in ecological communities worldwide

**DOI:** 10.1101/766790

**Authors:** Michael Kalyuzhny, Curtis H. Flather, Nadav M. Shnerb, Ronen Kadmon

**Affiliations:** Department of Ecology, Evolution and Behavior, Institute of Life Sciences, Hebrew University of Jerusalem, Givat-Ram, Jerusalem 91904, Israel.; USDA Forest Service, Rocky Mountain Research Station, Fort Collins, CO 80526, USA; Department of Physics, Bar-Ilan University, Ramat Gan 52900. Israel.

## Abstract

Ecological communities are assembled by colonization and extinction events, that may be regulated by ecological niches^1–5^. The most parsimonious explanation of local community assembly is the Dynamic Equilibrium (DE) model, which assumes that community dynamics is shaped by random colonization and extinctions events, effectively ignoring the effects of niches^1, 6^. Despite its empirical success in explaining diversity patterns^1, 5, 7^, it is unknown to what extent the assembly dynamics of communities around the globe are consistent with this model. Using a newly developed methodology, we show that in 4989 communities from 49 different datasets, representing multiple taxa, biomes and locations, changes in richness and composition are larger than expected by DE. All the fundamental assumptions of DE are violated, but the large changes in species richness and composition primarily stem from the synchrony in the dynamics of different species. These results indicate that temporal changes in communities are predominantly driven by shared responses of co-occurring species to environmental changes, rather than by inter-specific competition. This finding is in sharp contrast to the long-term recognition of competition as a primary driver of species assembly^8–12^. While ecological niches are often thought to stabilize species diversity and composition^4, 13, 14^, we found that they promote large changes in ecological communities.

## Main text

The primary goal of community ecology is understanding the processes shaping the dynamics and assembly of ecological communities. The most parsimonious view of ecological communities is that of dispersal assembly, postulating that communities are random assemblages of species that have arrived and have not yet gone extinct^15^. On top of that, niche assembly processes are viewed to act deterministically to shape community membership^3, 8^ and to stabilize species diversity^4^ and composition^13, 14^. A crucial question that has been investigated for decades is – what is the nature and magnitude of niche assembly processes?

Niche mechanisms are often detected as deviations from the predictions of dispersal assembly models^4, 8, 9, 16^. The simplest and most classical theory of dispersal assembly is the Dynamic Equilibrium model (DE), which assumes that diversity and composition are the result of stochastic colonization and extinction events of independent species, occurring at fixed rates^1, 6, 17^. DE predicts that species richness would fluctuate around an equilibrium, showing small temporal changes, while composition would show some degree of temporal turnover. Despite the central role that this model has played for decades and its empirical success in predicting static patterns^1, 5, 7^, its ability to explain ecological dynamics has rarely been quantitatively tested.

A few classical works have claimed to support the dynamic predictions of DE^18–20^. In recent years, following concerns over anthropogenic effects on ecological communities worldwide, there is growing interest in the dynamics of changes in diversity and composition. Several large-scale analyses have found that multiple communities around the globe show trends in richness^21–, 25^, seemingly violating the predictions of DE. Furthermore, some works have found compositional changes in excess of the predictions of DE and other null models in the vast majority of communities for which long-term records are available^21, 23^.

However, both classical and contemporary works have several important conceptual and methodological limitations. First, while the predictions of DE have been debated, very few works have tested its assumptions. This is unfortunate, since the assumptions are the mechanistic fundaments of any theory, and studying them can reveal the cause for observed deviations. Moreover, it is not clear to what degree the changes that have been found in richness and composition deviate from the expectations of DE. This is because richness trends are to be expected under DE if a community is initially not at equilibrium^6, 19^ and because the methods used to test compositional turnover have a very high chance of falsely detecting excessive changes when none exist^29, 30^.

Here we present the first rigorous, multi-taxa, global-scale evaluation of DE in 4989 communities from 49 different datasets (Fig S2). We ask three questions: (1) can temporal changes in richness and composition be explained by the DE model? (2) If not, what are the main deviations from the model predictions and, more importantly, the violation of which *assumptions* of the model are responsible for these deviations? (3) what ecological insights can we get from the observed deviations?

We reduce the high-dimensionality of community dynamics to five statistics, testing the assumptions and predictions of DE, using a newly developed methodology^29^. The *ds_rel_* and *dj* statistics quantify the magnitudes of changes in species richness and composition, respectively, beyond the expectation of DE. To examine the assumptions, we use *var_ex_* and *var_col_* to test whether extinction and colonization rates, respectively, are fixed, and the average covariance between species, 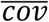, to test for species independence. The values and significance of these statistics are calculated for the time-series of composition over time in a community and are expected to be zero under DE. For more information see methods and ref. 29.

Our empirical analyses were done at two hierarchical levels –local community and dataset, whereas every community is a member of a dataset, and a dataset may include multiple relatively similar communities. We calculated the five statistics for every community, and then computed their average values for each of the 49 datasets. We present the results at the dataset level, and also at a single community level for the four largest datasets, all consisting of over 200 communities: the North Pacific Groundfish Observer (N.P. Fish), Konza Prairie LTER, the monitoring of fish in Skagerrak and the North American Breeding Bird Survey (N.A Birds).

We found widespread deviations from the predictions and assumptions of DE, both at the dataset and community resolution (Fig. 1). While DE predicts some temporal changes in richness, the observed changes were significantly higher (significant positive *ds_rel_*, Fig. 1a.) in 71.4% of datasets, and smaller changes then expected were never found. As for compositional turnover, we did find evidence for rates that are higher than DE expectations, but they occurred only in 30.6% of the datasets. (Fig. 1b). Testing the assumptions, we found that colonization and extinction rates were often not fixed, with periods of higher and periods of lower rates (mostly positive *var_ex_* and *var_col_* indicating that the variance in the length of presence and absence periods is higher than expected, Fig. 1c-d). Finally, while DE assumes species independence, covariance was significantly positive in 79.6% of the datasets, and never significantly negative (Fig. 1e). Qualitatively similar results were found at the level of single communities within datasets.

**Figure 1.**
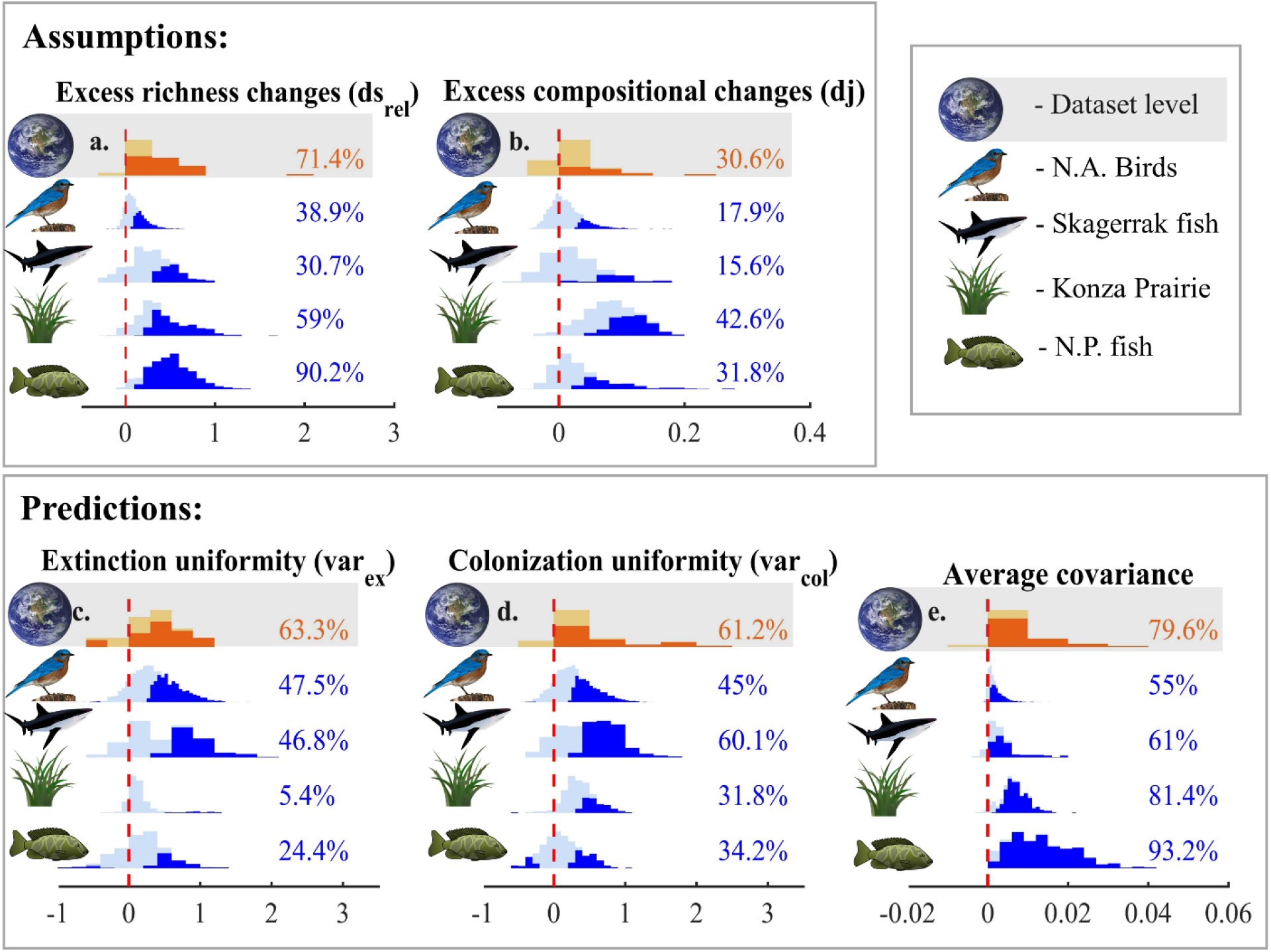
Histograms presenting the distributions of statistics quantifying the deviations from the predictions (limited changes in richness and composition, *ds_rel_* (a) and *dj* (b), respectively) and the assumptions (fixed extinction and colonization rates: *var_ex_* (c) and *var_col_* (d) respectively; independent species: average covariance (e)). In each panel, the upper histogram with gray background presents the dataset-level distribution, while the other histograms are at a community-level within the four largest datasets. Significant results are highlighted in the histogram with a darker color, and their percent of the total results is presented to the right. The expected value for all statistics is zero, marked with a red dashed line.

The larger changes then expected in richness and composition could arise from the non-uniformity of rates and/or from species dependence. To identify the cause, we regressed the statistics of deviations from the predictions (*ds_rel_* and *dj*) against the statistics of deviations from the assumptions. We found that *ds_rel_* is strongly correlated with 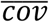 at the dataset and community levels (Table 1, Fig. 2), while the association with *var_ex_* and *var_col_* is much weaker, and often non-significant. Moreover, the effect of 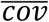 is considerably stronger for longer time series (Fig. 2, Table S1). Explaining the variation in excessive compositional changes proved to be more difficult, probably because of their small magnitude. While *dj* is not significantly associated with any explanatory variables at the dataset levels, for three of the large datasets, community-level *dj* is associated primarily with 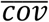 (Table 1). Similar results were obtained for more complex models (Table S1).

**Figure 2.**
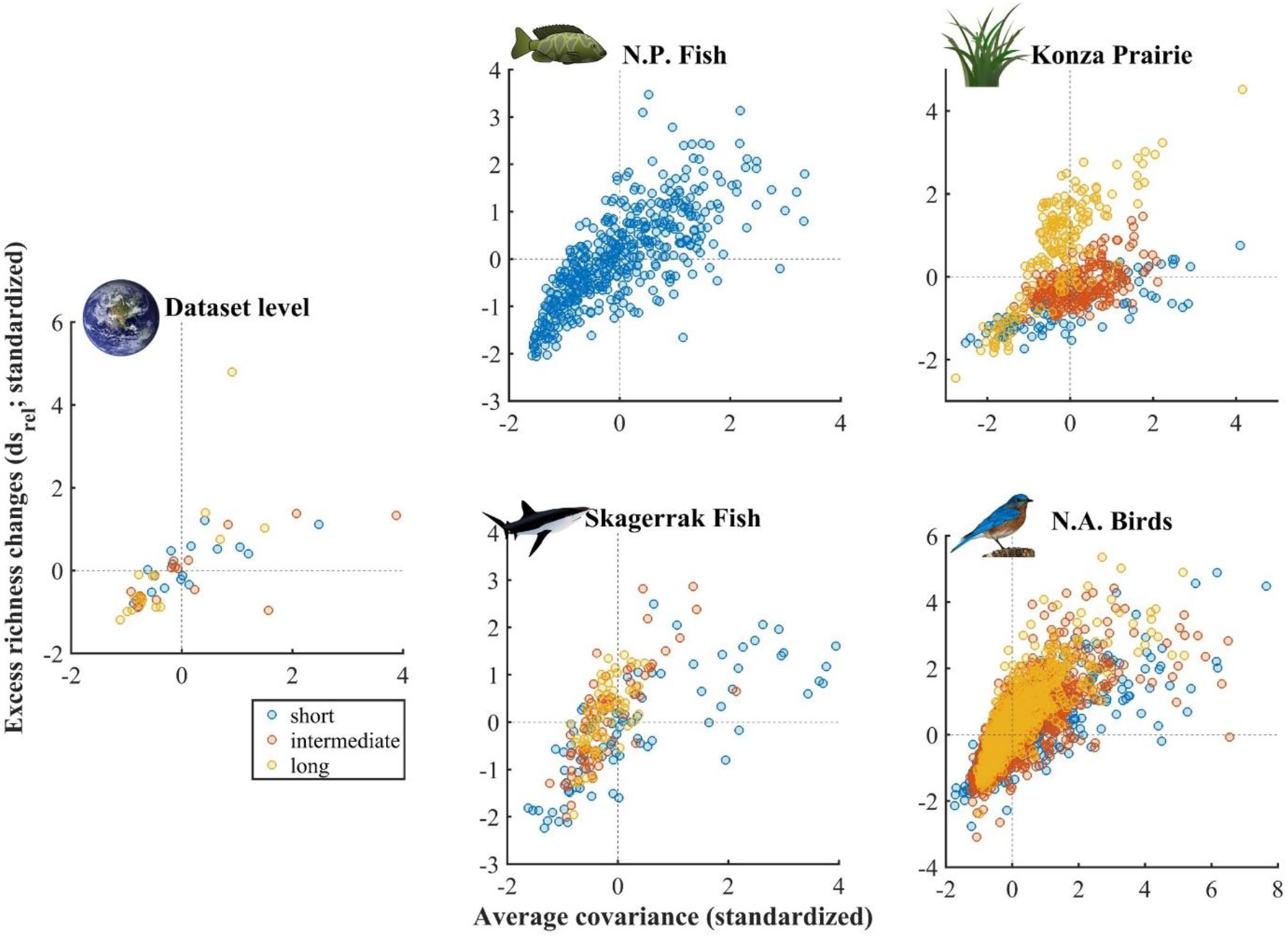
effect of average covariance on excessive changes in richness, *ds_rel_*, subdivided according to the length of the time series. Both variables are standardized. Results are presented at the dataset level (left, centered), as well as for single communities within the four largest datasets. Length categories are as follows: short time series are < 14 years for the dataset level analysis, < 20 years for Konza Prairie, < 30 for Skagerrak and < 15 for the N.A. Birds. Long time series are ≥ 25 years for the dataset level analysis, ≥ 30 years for Konza Prairie, ≥ 70 years for Skagerrak and ≥ 30 for the N.A birds. The N.P fish dataset is not divided because the lengths of it time series are 10-12 years.

**Table 1.** Regression models explaining the variation in *ds_rel_*, excessive richness changes, and *dj*, excessive compositional changes. The explanatory variables are the statistics quantifying deviations from DE assumptions - var_ex_ and var_col_, quantifying deviations from uniform rates extinction and colonization rates, respectively, and average covariance, var_col_. We present the standardized regression coefficients, the degrees of freedom of the errors and the coefficient of determination. *: P < 0.05, **: P < 0.01, ***: P < 0.001. Results are presented at the dataset level (in bold), and for single communities within the four largest datasets.

| *Standardized regression models for $ds_{rel}$ vs. assumptions* |  |  |  |  |  |
| --- | --- | --- | --- | --- | --- |
| *Coeff.* | **Dataset level** | N.P. Fish | Konza Prairie | Skagerrak Fish | N.A. Birds |
| $\overline{cov}$ | **0.686 \*\*\*** | 0.745 *** | 0.444 *** | 0.651 *** | 0.755 *** |
| $var_{ex}$ | **0.079** | 0.056 | -0.089 * | 0.174 ** | 0.077 *** |
| $var_{col}$ | **0.071** | -0.021 | 0.218 *** | 0.096 | 0.09 *** |
| DFE | **46** | 468 | 497 | 215 | 3530 |
| $R^2$ | **0.474** | 0.559 | 0.251 | 0.451 | 0.618 |
| *Standardized regression models for $dj$ vs. assumptions* |  |  |  |  |  |
| *Coeff.* | **Dataset level** | N.P. Fish | Konza Prairie | Skagerrak Fish | N.A. Birds |
| $\overline{cov}$ | **0.181** | 0.197 *** | 0.553 *** | 0.045 | 0.255 *** |
| $var_{ex}$ | **0.026** | 0.035 | 0.035 | -0.054 | 0.113 *** |
| $var_{col}$ | **0.126** | 0.045 | 0.177 *** | 0.083 | 0.06 ** |
| DFE | **46** | 468 | 497 | 215 | 3530 |
| $R^2$ | **0.057** | 0.046 | 0.322 | 0.007 | 0.098 |

Large changes in richness could be due to long-term trends or year to year variation (“noise”). To identify the nature of these changes, we quantified for each community the magnitude of noise and of the linear trend in richness. A regression of *ds_rel_* against these measures (Table 2) revealed that both trend and noise have a comparable contribution to the variation in richness changes (similar regression coefficients), but for different datasets their relative importance varies.

**Table 2.** Regression models explaining the variation in *ds_rel_*, excessive richness changes, using the statistics quantifying noise (SD of normalized year to year changes in richness) and trend (normalized Theil-Sen statistic), as well as their interactions with time series length. See methods for a detailed account of how these measures are calculated. We present the standardized regression coefficients, the degrees of freedom of the errors and the coefficient of determination. *: P < 0.05, **: P < 0.01, ***: P < 0.001. Both noise and trends contribute to large richness changes.

| *Standardized regression models for $ds_{rel}$ vs. trend and noise* |  |  |  |  |  |
| --- | --- | --- | --- | --- | --- |
| *Coeff.* | **Dataset level** | N.P. Fish | Konza Prairie | Skagerrak Fish | N.A. Birds |
| *length* | **0.564 \*\*\*** | 0.007 | 0.359 *** | 0.389 *** | 0.515 *** |
| *noise* | **0.685 \*\*\*** | 0.556 *** | 0.549 *** | 0.438 *** | 0.507 *** |
| *trend* | **0.426 \*\*\*** | 0.489 *** | 0.214 *** | 0.742 *** | 0.763 ** |
| *length\*noise* | **0.279 \*\*\*** | 0.098 *** | 0.165 *** | 0.058 | 0.052 *** |
| *length\*trend* | **0.128 \*\*\*** | 0.045 | 0.158 *** | 0.28 ** | 0.377 *** |
| *DFE* | **44** | 466 | 495 | 213 | 3528 |
| *R<sup>2</sup>* | **0.89** | 0.589 | 0.689 | 0.367 | 0.6 |

Not many generalities about the dynamics of ecological communities are known. To address this limitation, we have performed one of the most extensive tests ever performed in ecology of the ability of dynamic models to explain observed dynamics. We found that species richness indeed undergoes larger changes then expected under DE in multiple ecosystems worldwide. While previous works have focused primarily on trends, we have shown that year to year variation in richness (“noise”) has a similar strong contribution to excessive richness changes. Unlike previous studies using similar data^21^ only a minority of communities showed excessive compositional changes. For a comparison of the methodologies used, see ref. 29. While we expected that communities undergoing large richness changes would also show large compositional changes, this is not the case. We attribute this surprising result to the very large year-to-year compositional variation, with roughly 50% species turnover between consecutive years in many datasets. This is probably the result of extensive sampling errors across multiple datasets.

Our results shed light on the role of niche assembly beyond basic dispersal assembly processes. Both classical and contemporary studies of community assembly *in space* have revealed a prevalence of negative associations between species, with many pairs of species co-occurring less than expected by chance^8, 9, 11^. This has been interpreted as evidence for the possible importance of competition in shaping communities, along with other evidence, including experiments^12, 14^. However, while classical assembly studies tested whether static co-occurrence *patterns* in space resemble a random pattern, our approach is to examine the assembly *process* in action, testing if it resembles a random dispersal assembly process.

Given these results on negative associations, we expected that the arrival of one species would often increase the probability its competitors would go extinct, inducing negative average covariance between species. Surprisingly, this case is extremely rare in our data, where average covariance is almost exclusively positive. We interpret this as evidence that intra-specific competition is less important in shaping assembly dynamics (at least for species with intermediate abundance, for which changes are observed and are part of the analysis) then similar response of species to environmental changes (see appendix 1). This similar response to environmental changes, and not the non-uniformity of rates at the single species level, is the main driver of the large changes that are observed in richness and composition, as communities with higher covariance also have higher *ds_rel_* (and *dj*, to a lesser extent).

More work is needed to relate these excessive changes to specific drivers. Unlike previous works^21, 23^ we do not believe that such excessive changes can be necessarily interpreted as evidence for anthropogenic effects. This is because similar responses of species to the environment changes, which drive excessive changes, are generally expected in cases when the niches of species overlap, regardless of the causes of the environmental change.

Overall, we found that dispersal assembly alone is insufficient to explain the observed dynamics in ecological communities worldwide, and niche assembly processes play an important role. However, contrary to the expectation that niches buffer changes in composition and diversity^4, 13, 14^, we found that niches lead to synchrony in the dynamics of different species, which in turn leads to large temporal changes in richness and composition. This finding is in line with results on abundance changes^16, 26, 27^ and extinctions^28^ being driven by niches. We believe that in the face of the large pressure that ecological communities worldwide are undergoing, a synthetic view of the effect of ecological niches on community dynamics is crucial, and our work adds a piece to this puzzle.

## Acknowledgements

We wish to thank Micha Mandel for invaluable assistance during the development of the methodology, Eyal Ben-Hur for his considerably guidance on the graphical presentation of results, Shiran Faro for assembling the data and Nir Band for assistance with GIS. M.K. is supported by the Adams Fellowship Program of the Israel Academy of Sciences and Humanities.

## Methods

### Methodology for testing the Dynamic Equilibrium (DE) model

The properties of our newly developed methodology were studied in depth in ref. 29 of the main text. The methodology gets as an input a table of community composition (at a presence-absence level) over regular time intervals (without loss of generality we use the term “years”). The output is the values (effect size) and significance of a set of statistics that compare different aspects of community dynamics to the predictions and assumptions of DE. The two components of the approach are therefore the summary statistics and the null model to which they are compared. We use five statistics, one for each of the assumptions and predictions, and the Presence-Absence Redistribution wIthin periodS (PARIS) null model.

### The PARIS null model

The aim of the model is generating synthetic community time series of community composition that resemble the original time-series, but adhere to the assumptions of DE, namely, species independence and uniform colonization and extinction rates. This is done by the following algorithm, that is applied to single-species time series of presence-absence through time.

The PARIS algorithm preserves the initial state (presence or absence), the total number of years of presence and the total number of years of absence, the number of colonization and extinction events as well as the total number of periods of presence and periods of absence. The only thing that can change is the duration of these periods. Consider there were *y* years of presence in the empirical time series, separated into *s* periods by extinction events (that initiate periods of absence). The algorithm will choose *s* - 1 out of *y* - 1 years without replacement, simultaneously. Then, the events are assigned to happen after the chosen years. The analogous procedure is applied for periods of absence being separated by colonization events. If there was only one period of presence or absence, it will remain as in the original data in all resamples. Missing years are removed before the algorithm is applied and then are put back at their original timing.

For example, assume the original time series is [1 1 1 0 0 0 1 1 0 1], where “1” stands for a year of presence and “0” stands for a year of absence. There are six years of presence within three periods and four years of absence within two periods, so two extinction events and one colonization events should be assigned. For example, choosing extinction to occur after the 1^st^ and 2^nd^ years of presence and colonization to occur after the 2^nd^ year of absence will lead to the following synthetic time series: [1 0 0 1 0 0 1 1 1 1].

This algorithm generates independent time-series since it is applied to different species independently. Moreover, it generates events that are characteristic of a process with uniform rates (see ref. 29). Hence, it satisfies the assumptions of DE and can be used to generate multiple synthetic time series that “force” these assumptions on the data. An important property of the null model is that it preserves the initial state of the community, as well as the final state (the presence and absence of each species in the first and final year in the data).

### The statistics used to test DE

Each of the following statistics quantifies the deviation from DE in some aspect of the dynamics. They are constructed so that their mean if DE is true is zero.

To examine the magnitude of changes in richness, we use the statistic *ds_rel_*, which quantifies the normalized magnitude of temporal changes in species richness, beyond the expectation of DE:

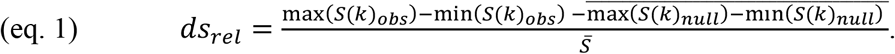

In other words, for the observed time series of species richness, *S*_*obs*_, we compute the change between the maximal richness and the minimal richness observed. The same change is then computed for every synthetic time-series generated by the null model, *S*_*null*_. Finally, the average average change under the null (the expected change) is subtracted from the observed change, and this is normalized by the average richness in the time series 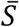 to ensure comparability between different studies.

Under the PARIS null model, richness in the first and last year in every randomization precisely equals the observed values. For this reason, it is useful to perform the calculation of *ds_rel_* for the “central” part of the time series, after “cutting” a proportion *k* of the years on both sides in the empirical data as well as in the randomizations. We used k = 0.06, so 0.06 of the total years (rounded up) will be “cut” on both sides for the calculation of *ds_rel_*. We performed a sensitivity analysis and found that using k = 0 would result in qualitatively similar (although weaker) results.

The statistic *dj* quantifies the magnitude of temporal changes in composition, beyond the expectation of DE:

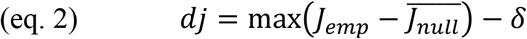

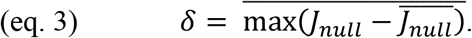

*J* is a curve describing the jaccard dissimilarity of each year w.r.t the first year. *J*_*emp*_ is this curve for the empirical data, while *J*_*null*_ is computed for a singe realization of the null model. Hence, eq. 2 computes the difference between *J*_*emp*_ and the expected *J*_*null*_ in the year when this difference is maximal. Since some deviations from the expected *J*_*null*_ will occur even under DE, we subtract the quantity *δ*, to ensure that dj will have a mean of 0 under DE. *δ* is the expected maximal deviation of a stochastic realization of the null model *J*_*null*_ from the expected *J*_*null*_.

To test the assumptions of uniformity in colonization and extinction rates, we use the statistics *var_col_* and *var_ex_*, respectively. These statistics are based on comparing the variance in the duration of absence and presence periods to the expectation under the null. For each species with two or more periods of presence or absence, we compute the variance in their durations (separately for presence and absence) and then standardize it using the null model. These standardized variances are then averaged over species:

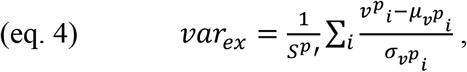

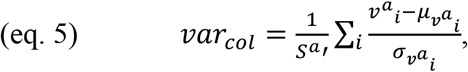

Where 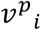 and 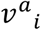 are the *observed* variances in the durations of presence and absence for species i, respectively; 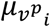 and 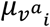 are the *expected* variances in the durations of presence and absence for species i, respectively; and 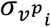 and 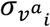 are the SD of the variances in the durations of presence and absence for species i, respectively, where the expectation and SD are calculated over resamples. These values are averaged over all species that have at least two events of presence or absence, respectively, *S*^*p*^′ and *S*^*a*^′.

To test species independence, we computed the covariance between every pair of species time-series, and averaged over the pairs.

All these statistics are expected to have a mean of zero under DE. A small exception is average covariance, which may have a positive value if the community initially is far from equilibrium. For all statistics except *dj*, a two-sided test is performed by comparing the observed statistics to a distribution of the statistic in 2·10^4^ randomizations and computing the p values. For *dj* the test is one sided.

At the dataset level, the values of the statistics are averaged over the communities within the dataset. For studies with > 200 communities, the significance of the deviation of this average from 0 is tested simply using a t test. For datasets with fewer communities, the randomization-produced statistics are recorded for each community separately. The test is performed by comparing the observed average statistic to the distribution of averaged (over communities) randomized statistics. For example, if a dataset consists of 10 communities, we first compute the average *dj* (or other statistic). We then compute and record 2·10^4^ randomization-generated *dj*s for each of the communities. Now we average the randomized statistics over the communities, to get 2·10^4^ average *dj*s, that will serve as the null distribution for the test.

### Properties of the methodology and interpretation

We have shown in ref 29. that the methodology generally has a type I error as predefined (α = 0.05 used in all our analyses). Moreover, it is reasonably robust to missing years, incomplete detection and false detection of species.

Several important comments regarding the interpretation of the output are in order. Since the PARIS null model fixes the initial state of the community, the methodology will not detect a deviation from DE if the community admits DE dynamics but the initial state is far from equilibrium. Furthermore, this methodology only considers species that are observed, and only those who have some colonization and extinction events contribute to the statistics. For this reason, the methodology will not detect cases when some species are never observed because of filtering (biotic or a-biotic), or the effect of very common or extremely rare species that never go extinct or are barely observed.

### Assembly of the database and data filtering

Our analyses are performed on tables of species by years, with information on presence and absence for each species at every year in a given site (community). For the analysis to make sense, we require that the data will be consistent in space (from a single site or locality), seasonality (every year the sampling is performed in the same months) and sampling effort. To achieve these goals, we assembled data from three sources: the newly published BioTIME database^1^, the Breeding Bird Survey (BBS)^2^ and further references that were available in previous publications^3, 4^ but were not included in the open version of BioTIME. A full list of these additional studies is included in table S3. A limited version of the BBS (300 sites) is included in the BioTIME but was discarded in favor of the full data (3533 sites included).

Following the recommendations of our methodological analysis^29^, we set the following quality criteria for all our data: we use data that has a timespan of at least 10 years, no more then 0.15 of which can be missing years (years when data is not available), and we required that at least 20 species have been observed sometime during the survey period.

In all cases, at most one community time-series is produced for a group of organisms (defined taxonomically or ecologically) in each site. Further details on the data gathering and cleaning are provided below according to the source of the data. The Matlab code for these analyses is included in the online appendix. A map presenting all datasets and communities is presented in Fig. S2.

### BioTIME

After downloading the database in CSV format (on 2/11/2018) We used the following algorithm to obtain time series that indeed satisfy our requirements. First, we discarded all studies with less then 10 years of time span and less than 20 species overall, all studies that had treatments and the BBS (study 195). next, for every study, we searched for unique sites. If data on plots was available, we treated every plot as a site and discarded every plot that had more then one latitude-longitude coordinate. If data on plots was not available (‘NA’), we considered every latitude-longitude coordinate as a site.

Some of the studies have no information on the season when sampling was performed. For these studies, we searched for the longest sequence of years that satisfies the quality conditions (timespan ≥ 10 years, ≤ 0.15 missing years, ≥ 20 species overall). If several such sequences were found with the same time span, we chose the sequence with the smallest number of missing years, and if several of these sequences had the same number of missing years, we (arbitrarily) chose the one with the earlier beginning. Sites without any time series satisfying our requirements were not used.

If data on sampling month was available, we wanted to make sure that the season of sampling is as consistent between years as possible. As a result, some records done in “a-typical” months, and some years lacking “typical” months would have to be discarded. Consequently, we had to balance between having time series with a longer time span in terms of years and having time series that include more months. This was achieved using the following algorithm.

We begin by constructing a binary table of years x months, indicating whether there is any data available for a given year-month combination. Then, the table is extended so that a year-month combination is coded as ‘available’ if one month before or after has any data. This is done to ignore small inconsistencies in sampling period. Next, we search for “good months” that are consistently sampled through the survey, that is, data for them is available for > *thresh_good_month* of the years. We used the value thresh_good_month = 0.9. Any year that has no data for a “good month” is discarded.

Next, we consider every one of the 12^2^ – 1 combinations of months. For each such combination, we find the number of months with available data, *m*, and the length of the ‘best’ sequence of years having these months, *y*. The ‘best’ sequence is found by finding the longest sequences satisfying the quality criteria (timespan ≥ 10 years, ≤ 0.15 missing years). Among them, the first among those with the fewest missing years is chosen. Now, the ‘value’ of each combination of months is calculated as *y*·*m^v^*, where *v* regulates the ‘value’ of a month w.r.t. a year. Since we need long sequences to have more power, we used v = 0.3. For example, this leads to 50 years with 5 months being better then 40 years with 10 months but worse than 40 years with 11 months. We find the combination of years and months that has the highest value and discard all data that does not belong to this combination. If no such combination with enough years, non-missing years and species is found, the entire site is discarded. This completes the selection of data for a single site.

Another issue that must be considered is that some sites have more then one ‘sample’ per site-year combination, and different years typically have different numbers of samples. In this case it is possible to subsample the data to the lowest number of samples per year (i.e. perform a rarefaction, as recommended by ref 1). However, this procedure may introduce some spatial variability (because the interpretation of ‘sample’ is different in different studies, e.g. in some studies it represents depth), or extra seasonality, complicating the interpretation of the results. Hence, while we used the rarefaction procedure in our preliminary analyses, getting qualitatively similar results, for our final analysis we discarded all sites that had more then one sample per year-site combination. The only exception was the Konza Prairie dataset, where this process would lead us to discard 92% out of the 500 sites in this unique dataset. Hence, for Konza Prairie we indeed rarefied the data.

Table S4 presents all the BioTIME datasets we included in the analysis.

### North American Breeding Bird Survey

The BBS is the largest single dataset of community dynamics in the world, yet it is notoriously famous for possible sampling errors^5–8^. In particular, changes in observer are known to sometimes xause different estimates^6, 7^. However, we could address this problem using the available information^2^ on the unique observer code of each observer.

The following algorithm was used for this task. Considering the records in a single site (BBS route), we search from the longest (all the data) to the shortest sequence of years for sequences satisfying the quality conditions (timespan ≥ 10 years, ≤ 0.15 missing years, ≥ 20 species overall), and where there is no evidence that changes in observer correspond to larger changes in richness or composition (the latter are quantified using Jaccard dissimilarity). To examine the effects of observer changes on changes in richness and composition, we use a one sided t test (with α = .05). We compare the magnitude of (absolute) changes in consecutive pairs of years when the observer changes to consecutive pairs when the observer remains the same. If this test is significant for either richness or compositional changes, this sequence of years is not used.

Once, for a given length of the time series, some satisfactory sequences are found, the algorithm picks the first (with the earlier start date) among the sequences with the least missing year. If no suitable sequence is found for a given length of time, the algorithm will look for shorter and shorter sequences sequentially, and if no suitable time series is found of the minimal length (10 years), the site will be discarded. The same years may therefore be considered in multiple runs of the algorithm searching for sequences, but eventually each BBS route will produce at most one time series.

### More datasets gathered

We found several references used in recent meta analyses^3, 4^ that were not included in the public version of the BioTIME. They were added to the database, and their details are available in table S2.

### Calculating normalized measures of noise and trends

Our aim was quantifying long-term trends and short-term fluctuations in richness, using normalized measures that would enhance comparison of communities with different richness.

To quantify trends at the community level we used the absolute value of the Theil-Sen statistic, normalized by the average richness in a community. The advantage of this statistic is that it is relatively robust to noise, making it inherently more independent of the noise measures. We took the absolute value because we were only interested in the magnitude of trends. At the dataset level we averaged this statistic over the communities that are members of the dataset.

The natural choice to quantify the magnitude of short-term richness fluctuations (“noise”) is the standard deviation of year-to-year richness changes:

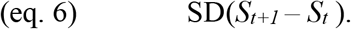

However, this statistic may require normalization, since it may depend on *S_t_*. In the population dynamics literature, it is known that if all the individuals in a population respond synchronously to environmental fluctuations, then changes in abundance would scale with initial abundance (a.k.a. environmental stochasticity). On the other hand, if individuals undergo birth and death independently, changes would scale with the square root of initial abundance (a.k.a. demographic stochasticity)^9, 10^. For a simple diffusion process, the magnitude of changes in the location of a random walker is independent of its location. We tried to classify our data roughly into these categories using the scheme suggested by ref. 11. For that aim, considering a set of richness time-series, we first computed the average change in every time series, 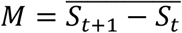. We then plotted the dependence of squared changes after subtracting the average, 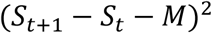 on *S*_*t*_ for all the time series in the set. Next, we estimated the regression equation

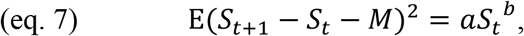

where E is the expectation operator (see results in table S3). If b ∼ 0, then richness changes do not scale with initial richness and there is no need to normalize richness changes (as in eq. 6). If b ∼ 1, richness changes should be normalized by √*S*_*t*_ (in analogy to demographic stochasticity), so the normalized measure for the magnitude of fluctuations would be

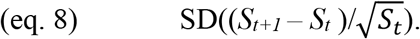

If b ∼ 2, richness changes should be normalized by *S_t_* (in analogy to environmental stochasticity), so the normalized measure for the magnitude of richness changes would be:

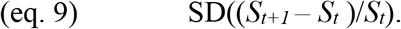

The analysis was performed at the dataset level as well as at the community level for the four largest datasets. At the dataset level, we pooled together all the richness changes from all the communities except for the BBS (since these communities strongly affected the result), and estimated *b*. Then, for each of the four largest datasets, we separately pooled the richness changes and performed the regression to estimate *b*.

We found that for all the communities together, b ∼ 1. Hence, for the dataset level analysis, we used eq. 8 as the measure for the magnitude of year to year fluctuations in richness in each community. These values were then averaged over the communities that are members of the dataset. For the community level analysis, we found b only a little larger than 0 for all datasets but the Konza Prairie, where b was close to two. Hence, we used eq. 9 to quantify the magnitude of richness changes for the communities within the Konza Prairie dataset, and eq. 6 for communities in the other datasets. See table S3 for the results of the fits of eq. 7.

While estimating fluctuation scaling is challenged by deterministic effects (part of the changes are due to the deterministic dynamics), sampling errors etc., so our results are only suggestive, we believe that our attempts to use different normalizations, getting similar results, can be considered an effective robustness check.

## Supplementary materials

### Appendix 1: interpreting covariance patterns between species

One of our main results was the predominance of positive average covariance in the presence-absence time series in ecological communities across the globe. We further found that changes in richness and composition are largely determined by the level of this covariance. Here we theoretically investigate the possible mechanistic drivers of this pattern.

Negative average temporal covariance between species can be depicted as species replacing each other through time, with the arrival of some corresponding to an increase in the extinction of others. Our intuition suggests that such negative covariance is the result of competitive interactions or different responses to environmental changes. On the other hand, positive covariance or synchrony can be viewed as species arriving and going extinct simultaneously, with the presence of some species increasing the probability others will be present. We believe this could stem from facilitative interactions or similar response to environmental changes. Consequently, we suggest that the overall covariance in a community can indicate which are the dominant process in determining the observed patterns of colonization and extinction, and, accordingly, changes in richness and composition.

Most of the datasets we studied consist of relatively similar species in terms of trophic level, taxon and size, making us believe that the interactions among them are primarily competitive. Therefore, the net positive effect that we found indicates that competition and different responses to environmental changes (causing negative covariance) are less important than similar response to environmental changes (causing positive covariance) in shaping observed community dynamics. This intuition should of course be tested. For that aim, we used simulations of population dynamics models that incorporate competition, immigration and temporal environmental vitiation and covariation. Our aim was to study how the strength of competitive interactions and the degree of similarity in the response to environmental variation affect the average covariance in presence-absence time series.

We ran simulations of a stochastic, discrete-time multispecies Ricker model^1, 2^. In this model, the expected population of species *i* at time *t*+1, *N*_*i*,*t*+1_, is:

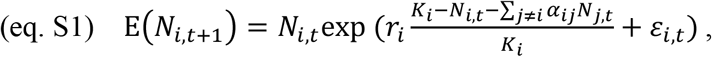

where *r*_*i*_ and *K*_*i*_ are the growth rate and carrying capacity of species *i*, respectively, *α*_*ij*_ is the per capita effect of an individual of species *j* on the growth of species *i*, representing inter-specific interactions, and *ε*_*i*,*t*_ represents stochastic fluctuations in the growth rate due to environmental changes. *ε*_*i*,*t*_ is normally distributed with a mean of 0, variance of *σ*_*e*_^2^ and the covariance between every pair of species is assumed to be *C*, representing the degree to which species respond similarly to environmental variation.

While eq. S1 represents the expected population of species i at time t, the actual population size is drawn from a Poisson distribution: *N*_*i*,*t*+1_∼Poisson(E(*N*_*i*,*t*+1_)). This introduces demographic stochasticity, that is, random variation between individuals in demography, as well as the discreteness of individual, which allows species to go stochastically extinct. Finally, after the local demography step described above, we introduce a single immigrant per time step, who is chosen uniformly from the *S_reg_* species available in the species pool.

The simulations were initialized with one individual of each of the species in the pool. Then they were run for 10^5^ time steps, and the first 2·10^3^ time steps were considered equilibration time and removed. The covariance matrix of the presence-absence time series was then computed. This allowed us to calculate the average of the off-diagonal elements, representing the covariance between pairs of species, averaged over pairs. This is precisely the statistic we consider in the main text.

We present in Fig. S1 results for the following parameters. We assumed that the pool consists of 20 species, each with r_i_ = 0.1 and K = 1000. The *α*_*ij*_ s are drawn from a normal distribution with mean 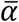 and SD of *σ*_*α*_, and negative values are transformed into positive ones by taking their absolute values. This results in purely competitive dynamics. We took *σ*_*α*_ to be 0.05 and considered values of 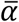 between 0 and 0.5 (in steps of 0.05), controlling the magnitude of competitive interactions. We considered *σ*_*e*_^2^ = 0.3, and C to be 0 to 0.3 (in steps of 0.05). The extreme values, C = 0 and C = *σ*_*e*_^2^, represent scenarios of fully independent and full synchronous responses to environmental variation, respectively.

We found that when competitive interactions are strong and environmental covariance is low, the average covariance in presence-absence is negative, albeit only slightly lower than zero (Fig S1). This is to be expected, since, mathematically, there are tight limits on how negatively a set of random variables can be correlated. On the other hand, when interactions are weak and environmental covariation is high, average covariance tends to be positive. Other parameters are in the middle between these cases, with more covariation and less competition generally leading to more positive covariance in presence-absence. These qualitative conclusions also hold in other parameter regimes we examined.

These results lead us to believe that the positive covariance we found in the data is indeed the signature of similar response of species to environmental variation overwhelming competition in generating community covariance patterns. This does not imply, however, that competitive interactions are weak in absolute terms, and they can still strongly affect overall richness and composition. Yet, our results demonstrate that the changes in richness and composition, that are associated with average covariance, are primarily governed by similar responses to environmental changes. While we examined a very specific model, recent works have shown for a similar model (continuous-time Lotka-Volterra) that many “macroscopic” patterns of communities are fully determined by a few parameters. In particular, these parameters are the variances of intra-specific and inter-specific interactions, the mean strength of inter-specific interactions, and their symmetry (does i affect j in the same manner j affects i?)^3, 4^, which are analogous to many of the parameters that we have considered. Moreover, these works have shown that many more complex models essentially map into this simple model for a single trophic level. This makes us believe that despite the simplicity of our model, the qualitative results can be generalized for many ecological communities.

**Figure S1.**
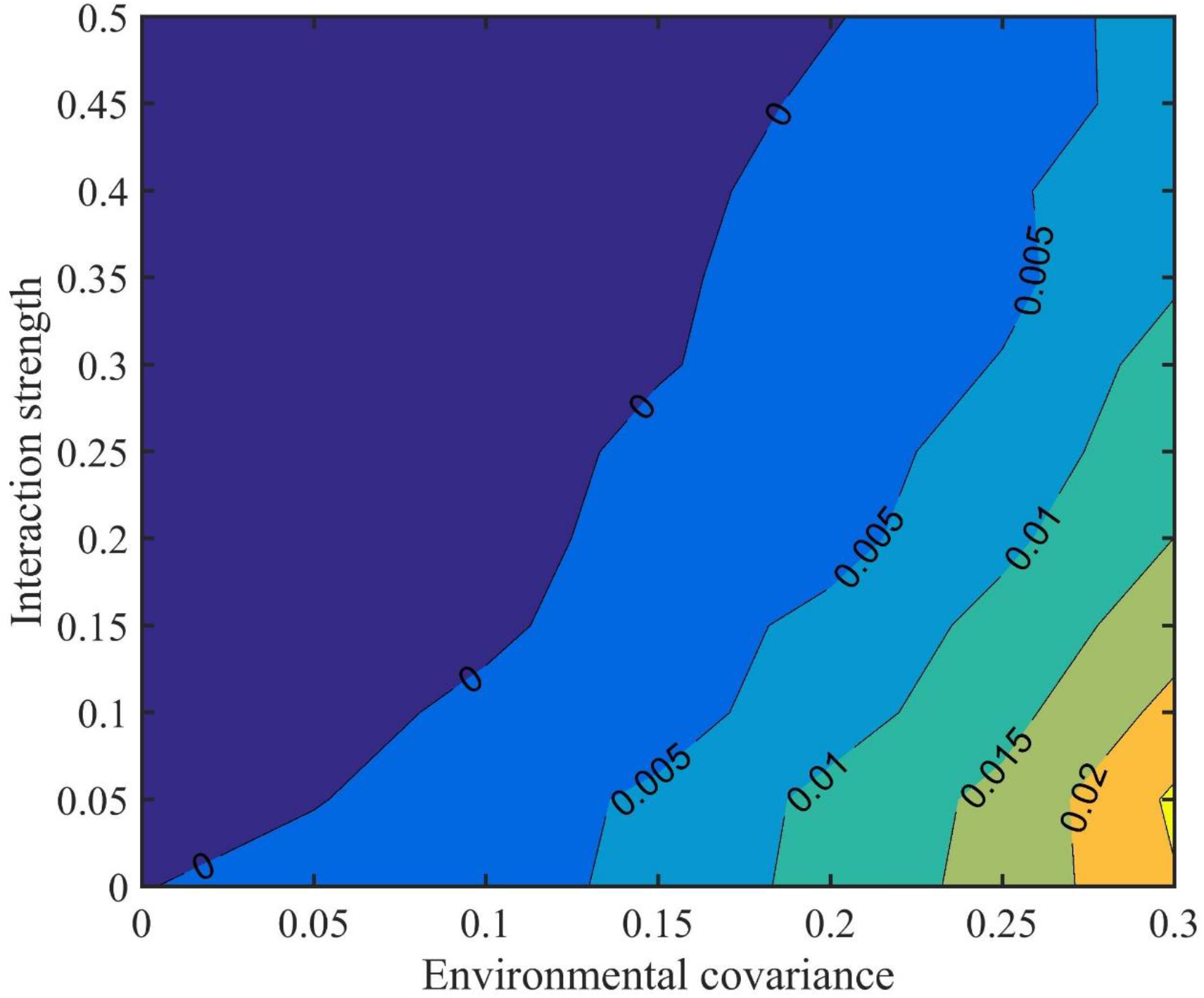
Contour plot of the combined effect of mean interactions strength (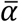, Y axis) and environmental covariance (C, X axis) on the average covariance in the presence-absence time series of species.

## Supplementary figures and tables

**Table S1.** More complex regression models explaining the variation in *ds_rel_*, excessive richness changes, and *dj*, excessive compositional changes, using the statistics quantifying deviations from DE assumptions. These models include interactions between average covariance and rate non-uniformity statistics as well as the length of the time series and its interaction with average covariance. We present the standardized regression coefficients, the degrees of freedom of the errors and the coefficient of determination. *: P < 0.05, **: P < 0.01, ***: P < 0.001. Results are presented at the dataset level (in bold), as well as for single communities within the four largest datasets. *ds_rel_* is strongly and positively associated with covariance, and much less with the rate-uniformity statistics. *dj* is less well explained, but for the single studies it is associated with covariance as well.

| Standardized regression models for $ds_{rel}$ vs. assumptions | | | | | |
| --- | --- | --- | --- | --- | --- |
| Coeff. | Dataset level | N.P. Fish | Konza Prairie | Skagerrak Fish | N.A. Birds |
| $\overline{cov}$ | **0.842 \*\*\*** | 0.777 *** | 0.542 *** | 1.044 *** | 0.86 *** |
| $var_{ex}$ | **-0.238** | 0.029 | -0.078 * | 0.089 | -0.037 ** |
| $var_{col}$ | **-0.243** | -0.034 | -0.067 * | -0.013 | -0.034 * |
| Length | **0.515 \*\*** | 0.052 | 0.718 *** | 0.303 *** | 0.243 *** |
| $\overline{cov}^* var_{ex}$ | **0.007** | 0.063 * | 0.036 | -0.021 | -0.119 *** |
| $\overline{cov}^* var_{col}$ | **-0.168** | -0.007 | -0.064 * | -0.096 | -0.101 *** |
| $\overline{cov}^* length$ | **0.557 \*\*** | 0.106 *** | 0.292 *** | 0.507 *** | 0.312 *** |
| DFE | **42** | 464 | 493 | 211 | 3526 |
| $R^2$ | **0.726** | 0.586 | 0.717 | 0.527 | 0.672 |
| Standardized regression models for $dj$ vs. assumptions | | | | | |
| Coeff. | Dataset level | N.P. Fish | Konza Prairie | Skagerrak Fish | N.A. Birds |
| $\overline{cov}$ | **0.26** | 0.21 *** | 0.559 *** | 0.169 | 0.273 *** |
| $var_{ex}$ | **-0.135** | 0.036 | -0.053 | -0.03 | 0.059 ** |
| $var_{col}$ | **-0.046** | 0.051 | 0.098 * | 0.103 | 0.011 |
| Length | **0.305** | 0.003 | 0.134 ** | -0.027 | 0.114<br>*** |
| $\overline{\text{cov}}^*$<br>$\text{var}_{\text{ex}}$ | **-0.013** | -0.017 | -0.084 * | 0.134 | 0.011 |
| $\overline{\text{cov}}^*$<br>$\text{var}_{\text{col}}$ | **-0.1** | -0.036 | -0.035 | -0.149 | -0.11 *** |
| $\overline{\text{cov}}^* \text{le}$<br>ngth | **0.189** | 0.059 | -0.023 | 0.14 | 0.014 |
| DFE | **42** | 464 | 493 | 211 | 3526 |
| $R^2$ | **0.103** | 0.053 | 0.348 | 0.033 | 0.112 |

**Table S2.** List of datasets gathered from the references of previous meta-analyses that were not included in the open version of BioTIME.

| Study name | Ref. | Link | Organisms | Communities | Comments | Lat | Lon |
| --- | --- | --- | --- | --- | --- | --- | --- |
| The long-term study of the fish and crustacean community of the Bristol Channel | 5 | http://www.pisces-conservation.com/ | Fish | 1 | Only data on fish used | 51.2 | -3.1 |
| Jornada experimental range – plant cover | 6 | https://jornada.nmsu.edu/content/jornada-experimental-range-permanent-quadrat-chart-data-beginning-1915-plant-cover | Plants | 1 |  | 32.500 | -106.80 |
| Jornada experimental range – plant density | 6 | https://jornada.nmsu.edu/content/jornada-experimental-range-permanent-quadrat-chart-data-beginning-1915-plant-density | Plants | 1 |  | 32.500 | -106.80 |
| Eastern Wood Breeding Bird Data | 7 | https://onlinelibrary.wiley.com/doi/pdf/10.1002/9780470999592.app2 | Birds | 1 | Years when some species were not counted were discarded | 51.2964 | -0.3835 |
| Long-term monitoring of mammals in the face of biotic and abiotic influences at a semiarid site in north-central Chile | 8 | http://www.esapubs.org/archive/ecol/E094/084/metadata.php | Mammals | 1 | Merged to one community table | -30.681 | -71.66 |
| Long-Term Data from Fields Recovering after Sugarcane, | 9 | https://www.hindawi.com/journals/dpis/2013/468973/ | Plants | 6 |  | 0.03 | -78.5 |
| Banana, and Pasture Cultivation in Ecuador |  |  |  |  |  |  |  |
| Thirty years of permanent vegetation plots, Mount St. Helens, Washington | 10 | http://esapubs.org/archive/ecol/E091/152/metadata.htm | Plants | 1 |  | 46.220 | -122.20 |
| Species interactions during succession in the western cascade range of Oregon, 1990 to present | 11 | http://andlter.forestry.oregonstate.edu/data/abstract.aspx?dbcode=TP103 | Plants | 1 |  | 44.174 | -122.25 |

**Table S3.** Results of the fits of squared changes in species richness (after removing the average change) vs. initial richness. The results are presented for all the data pulled together, excluding the BBS, as well as for the data of four large datasets pulled together. 95% confidence intervals for the power, b, are presented along with the coefficient of determination of the model.

|  | All data<br>excluding BBS | NP Fish | Konza Prairie | Skagerrak Fish | BBS |
| --- | --- | --- | --- | --- | --- |
| b in model<br>$E(S_{t+1} - S_t - M)^2 = aS_t^b$ | $1.106 \pm 0.042$ | $-0.1085 \pm 0.1010$ | $1.705 \pm 0.088$ | $0.2609 \pm 0.0841$ | $0.2144 \pm 0.0452$ |
| $R^2$ of $y = a \cdot x^b$ | 0.0975 | 0.0006 | 0.0727 | 0.0021 | 0.0012 |

**Figure S2.**
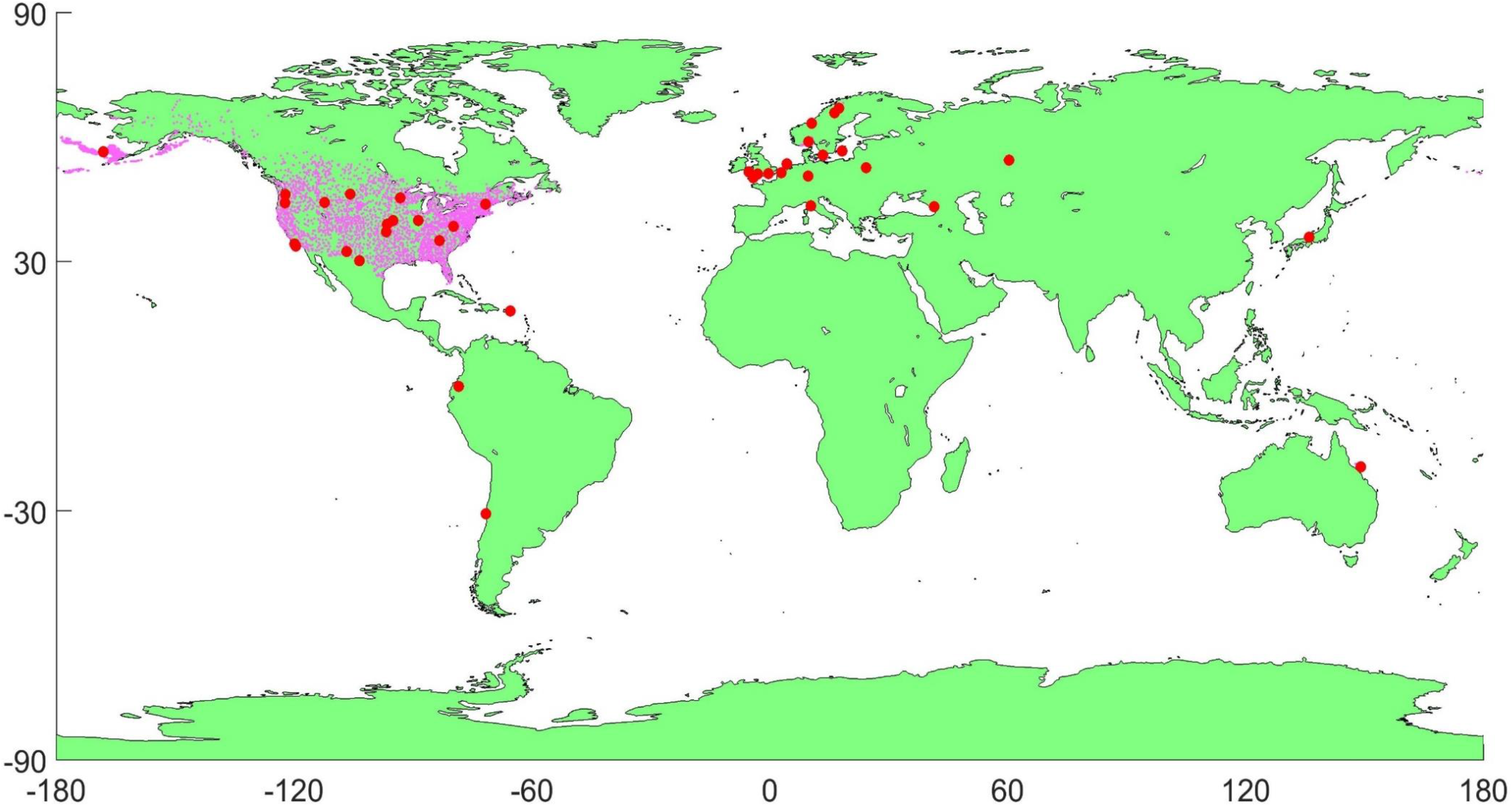
map of all communities (small purple dots) and datasets (red circles) use in our analysis. For the datasets, the central coordinate is presented.

**Table S4.**
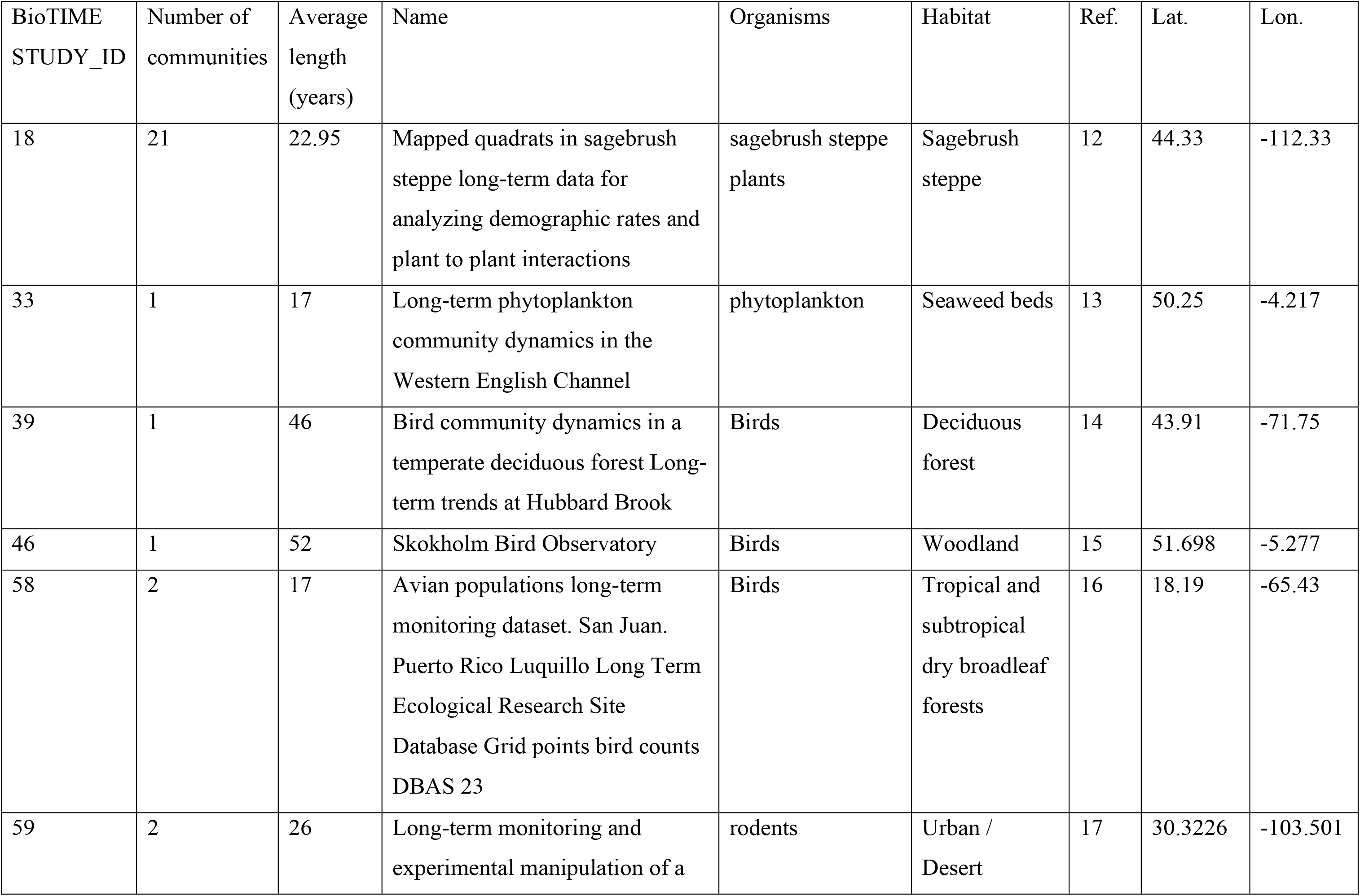
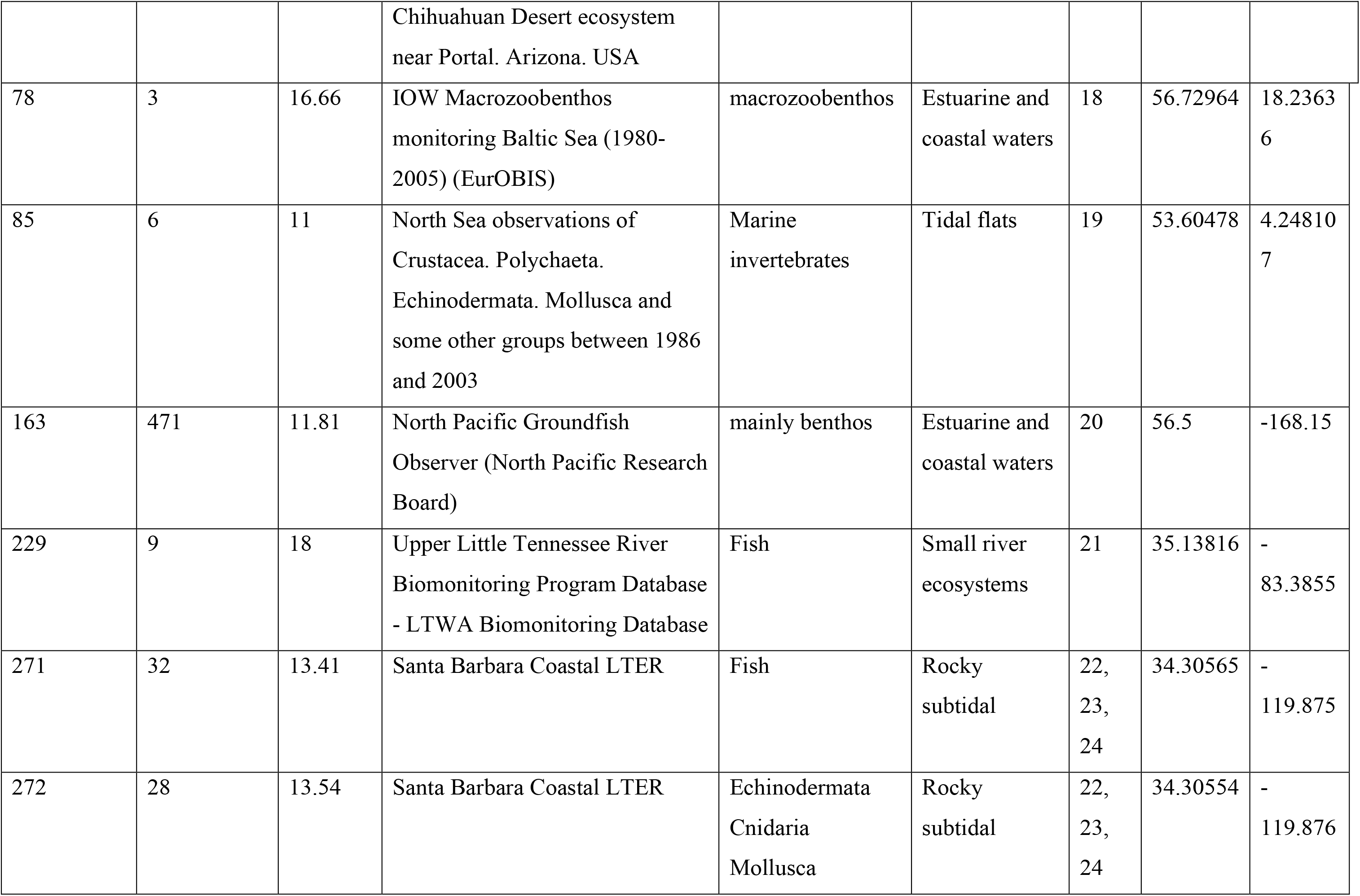
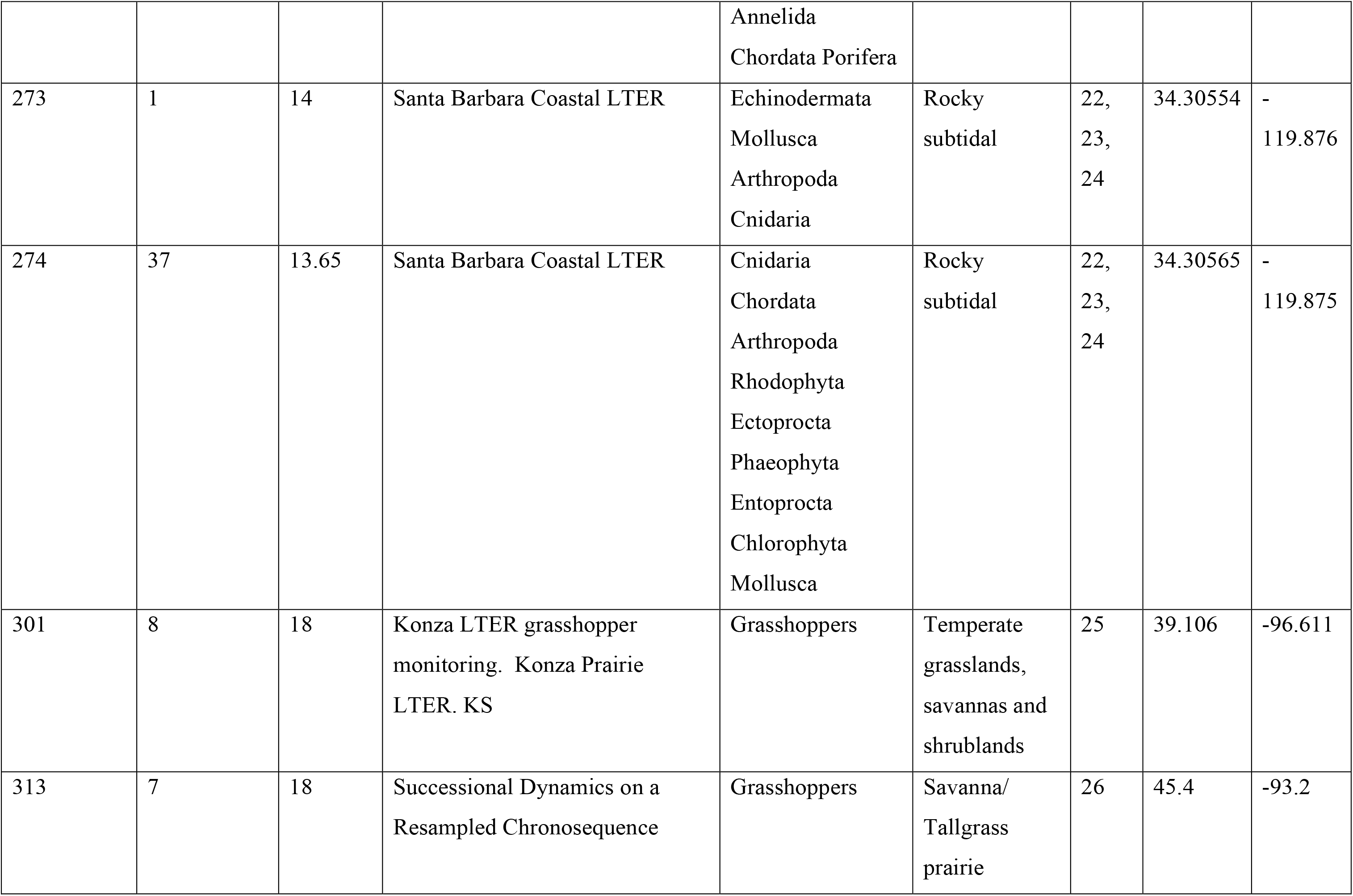
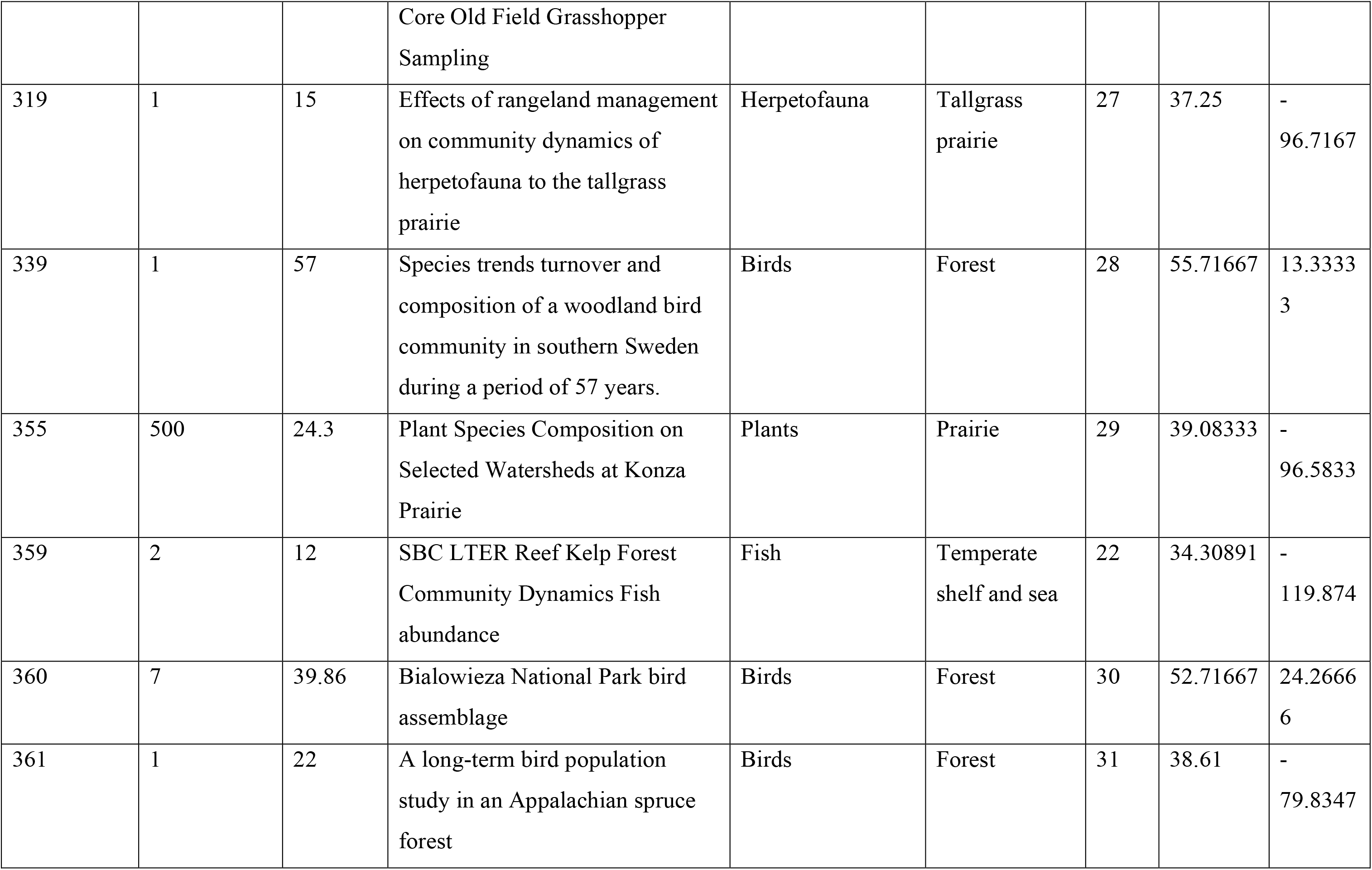
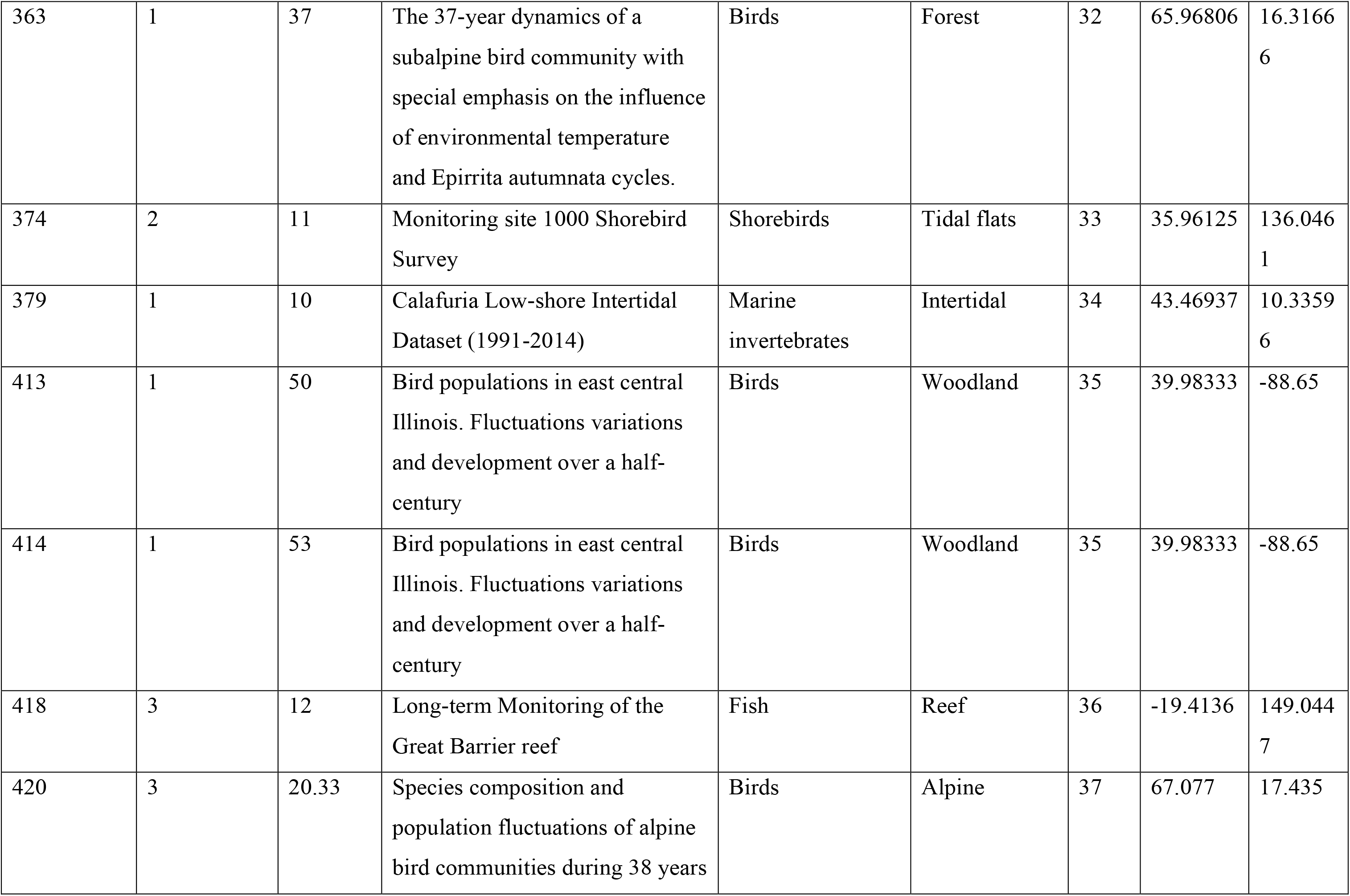
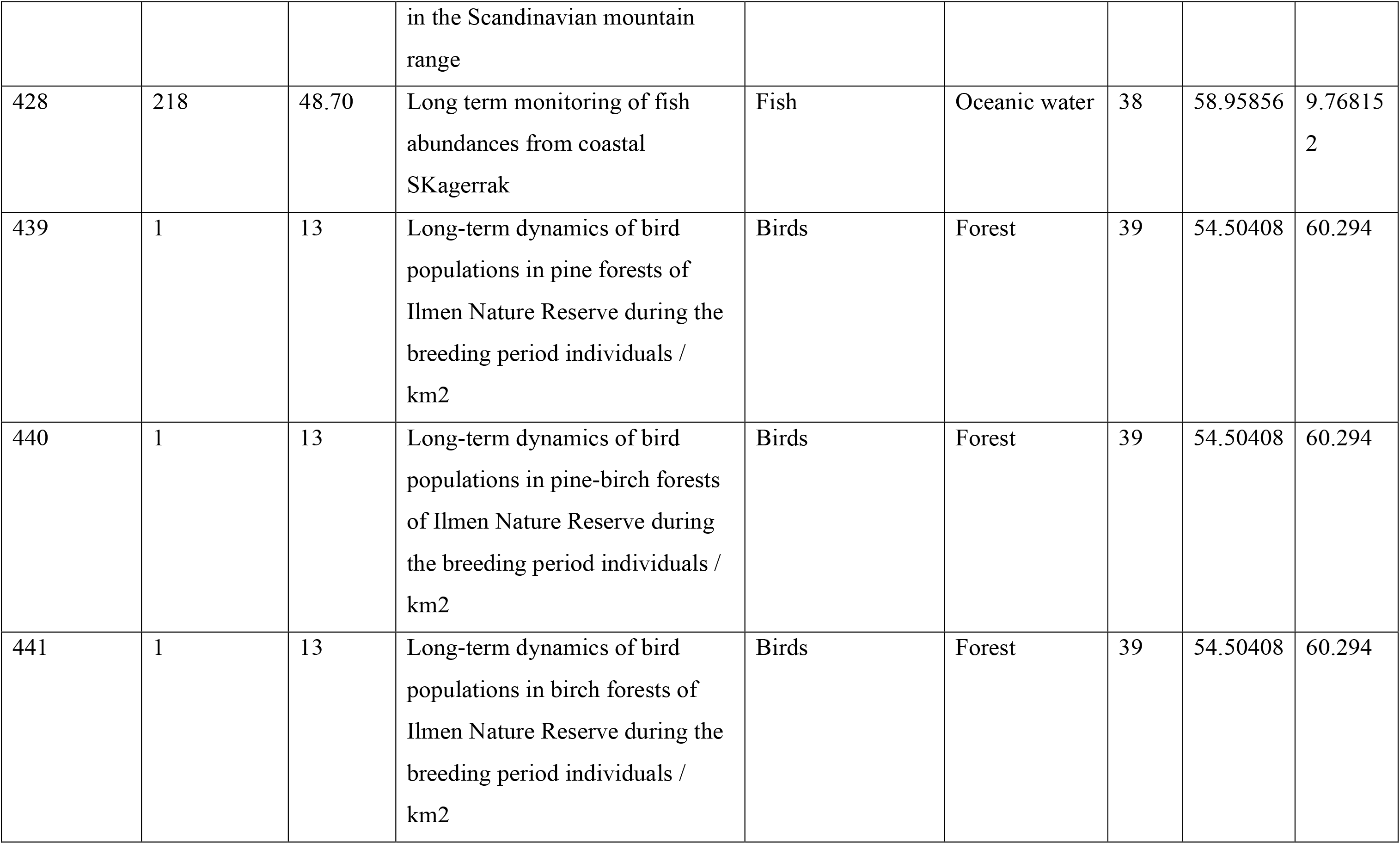
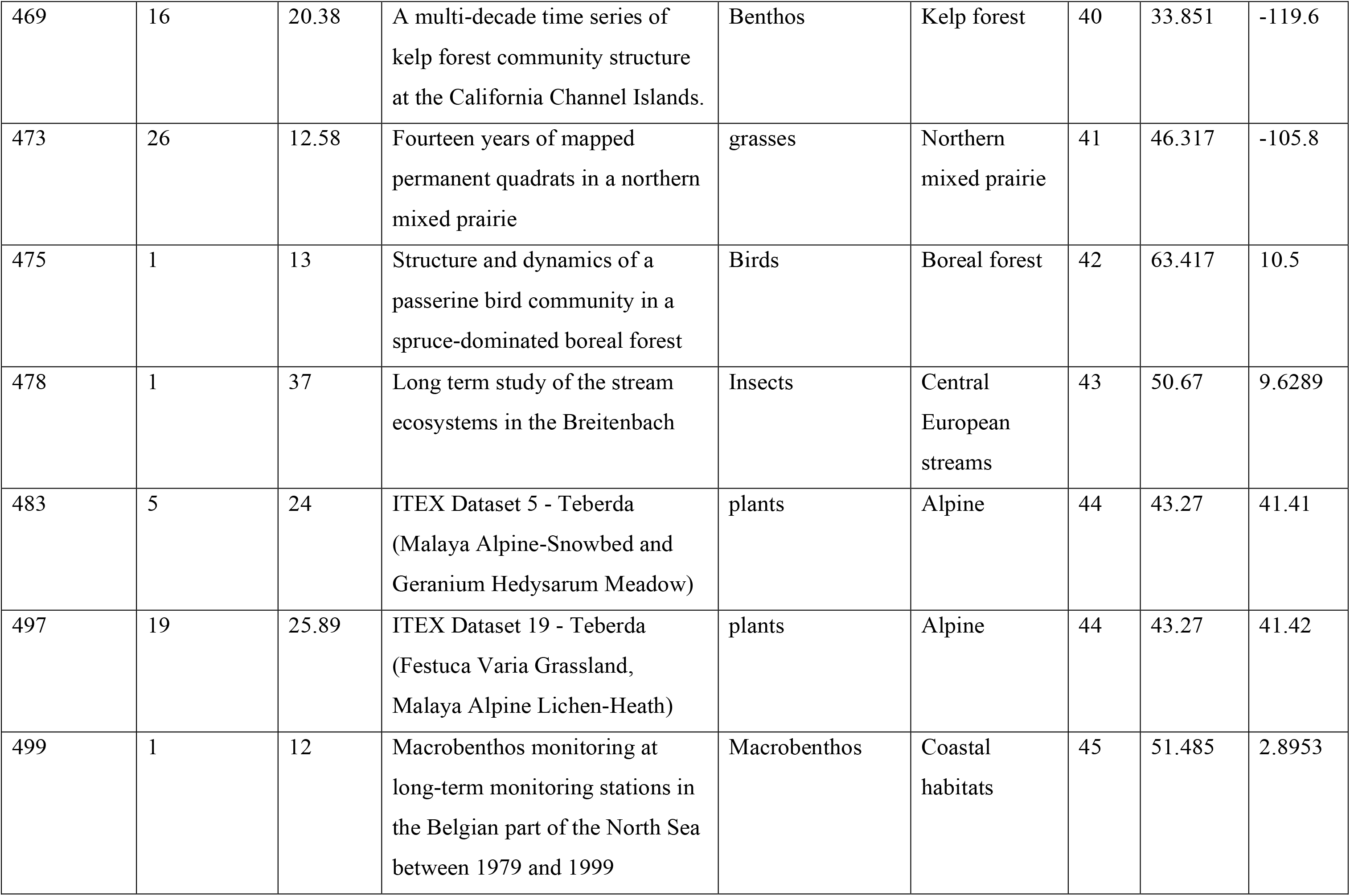
the datasets from BioTIME used for the analysis, along with the number of communities they consist of and their average timespan.

| BioTIME<br>STUDY_ID | Number of<br>communities | Average<br>length<br>(years) | Name | Organisms | Habitat | Ref. | Lat. | Lon. |
| --- | --- | --- | --- | --- | --- | --- | --- | --- |
| 18 | 21 | 22.95 | Mapped quadrats in sagebrush steppe long-term data for analyzing demographic rates and plant to plant interactions | sagebrush steppe plants | Sagebrush steppe | 12 | 44.33 | -112.33 |
| 33 | 1 | 17 | Long-term phytoplankton community dynamics in the Western English Channel | phytoplankton | Seaweed beds | 13 | 50.25 | -4.217 |
| 39 | 1 | 46 | Bird community dynamics in a temperate deciduous forest Long-term trends at Hubbard Brook | Birds | Deciduous forest | 14 | 43.91 | -71.75 |
| 46 | 1 | 52 | Skokholm Bird Observatory | Birds | Woodland | 15 | 51.698 | -5.277 |
| 58 | 2 | 17 | Avian populations long-term monitoring dataset. San Juan. Puerto Rico Luquillo Long Term Ecological Research Site Database Grid points bird counts DBAS 23 | Birds | Tropical and subtropical dry broadleaf forests | 16 | 18.19 | -65.43 |
| 59 | 2 | 26 | Long-term monitoring and experimental manipulation of a | rodents | Urban / Desert | 17 | 30.3226 | -103.501 |
|  |  |  | Chihuahuan Desert ecosystem<br>near Portal. Arizona. USA |  |  |  |  |  |
| 78 | 3 | 16.66 | IOW Macrozoobenthos<br>monitoring Baltic Sea (1980-2005) (EurOBIS) | macrozoobenthos | Estuarine and<br>coastal waters | 18 | 56.72964 | 18.2363<br>6 |
| 85 | 6 | 11 | North Sea observations of<br>Crustacea. Polychaeta.<br>Echinodermata. Mollusca and<br>some other groups between 1986<br>and 2003 | Marine<br>invertebrates | Tidal flats | 19 | 53.60478 | 4.24810<br>7 |
| 163 | 471 | 11.81 | North Pacific Groundfish<br>Observer (North Pacific Research<br>Board) | mainly benthos | Estuarine and<br>coastal waters | 20 | 56.5 | -168.15 |
| 229 | 9 | 18 | Upper Little Tennessee River<br>Biomonitoring Program Database<br>- LTWA Biomonitoring Database | Fish | Small river<br>ecosystems | 21 | 35.13816 | -<br>83.3855 |
| 271 | 32 | 13.41 | Santa Barbara Coastal LTER | Fish | Rocky<br>subtidal | 22,<br>23,<br>24 | 34.30565 | -<br>119.875 |
| 272 | 28 | 13.54 | Santa Barbara Coastal LTER | Echinodermata<br>Cnidaria<br>Mollusca | Rocky<br>subtidal | 22,<br>23,<br>24 | 34.30554 | -<br>119.876 |
|  |  |  |  | Annelida<br>Chordata Porifera |  |  |  |  |
| 273 | 1 | 14 | Santa Barbara Coastal LTER | Echinodermata<br>Mollusca<br>Arthropoda<br>Cnidaria | Rocky<br>subtidal | 22,<br>23,<br>24 | 34.30554 | -<br>119.876 |
| 274 | 37 | 13.65 | Santa Barbara Coastal LTER | Cnidaria<br>Chordata<br>Arthropoda<br>Rhodophyta<br>Ectoprocta<br>Phaeophyta<br>Entoprocta<br>Chlorophyta<br>Mollusca | Rocky<br>subtidal | 22,<br>23,<br>24 | 34.30565 | -<br>119.875 |
| 301 | 8 | 18 | Konza LTER grasshopper<br>monitoring. Konza Prairie<br>LTER. KS | Grasshoppers | Temperate<br>grasslands,<br>savannas and<br>shrublands | 25 | 39.106 | -96.611 |
| 313 | 7 | 18 | Successional Dynamics on a<br>Resampled Chronosequence | Grasshoppers | Savanna/<br>Tallgrass<br>prairie | 26 | 45.4 | -93.2 |
|  |  |  | Core Old Field Grasshopper Sampling |  |  |  |  |  |
| 319 | 1 | 15 | Effects of rangeland management on community dynamics of herpetofauna to the tallgrass prairie | Herpetofauna | Tallgrass prairie | 27 | 37.25 | - 96.7167 |
| 339 | 1 | 57 | Species trends turnover and composition of a woodland bird community in southern Sweden during a period of 57 years. | Birds | Forest | 28 | 55.71667 | 13.3333 3 |
| 355 | 500 | 24.3 | Plant Species Composition on Selected Watersheds at Konza Prairie | Plants | Prairie | 29 | 39.08333 | - 96.5833 |
| 359 | 2 | 12 | SBC LTER Reef Kelp Forest Community Dynamics Fish abundance | Fish | Temperate shelf and sea | 22 | 34.30891 | - 119.874 |
| 360 | 7 | 39.86 | Bialowieza National Park bird assemblage | Birds | Forest | 30 | 52.71667 | 24.2666 6 |
| 361 | 1 | 22 | A long-term bird population study in an Appalachian spruce forest | Birds | Forest | 31 | 38.61 | - 79.8347 |
| 363 | 1 | 37 | The 37-year dynamics of a subalpine bird community with special emphasis on the influence of environmental temperature and *Epirrita autumnata* cycles. | Birds | Forest | 32 | 65.96806 | 16.31666 |
| 374 | 2 | 11 | Monitoring site 1000 Shorebird Survey | Shorebirds | Tidal flats | 33 | 35.96125 | 136.0461 |
| 379 | 1 | 10 | Calafuria Low-shore Intertidal Dataset (1991-2014) | Marine invertebrates | Intertidal | 34 | 43.46937 | 10.33596 |
| 413 | 1 | 50 | Bird populations in east central Illinois. Fluctuations variations and development over a half-century | Birds | Woodland | 35 | 39.98333 | -88.65 |
| 414 | 1 | 53 | Bird populations in east central Illinois. Fluctuations variations and development over a half-century | Birds | Woodland | 35 | 39.98333 | -88.65 |
| 418 | 3 | 12 | Long-term Monitoring of the Great Barrier reef | Fish | Reef | 36 | -19.4136 | 149.0447 |
| 420 | 3 | 20.33 | Species composition and population fluctuations of alpine bird communities during 38 years in the Scandinavian mountain range | Birds | Alpine | 37 | 67.077 | 17.435 |
| 428 | 218 | 48.70 | Long term monitoring of fish abundances from coastal SKagerrak | Fish | Oceanic water | 38 | 58.95856 | 9.768152 |
| 439 | 1 | 13 | Long-term dynamics of bird populations in pine forests of Ilmen Nature Reserve during the breeding period individuals / km2 | Birds | Forest | 39 | 54.50408 | 60.294 |
| 440 | 1 | 13 | Long-term dynamics of bird populations in pine-birch forests of Ilmen Nature Reserve during the breeding period individuals / km2 | Birds | Forest | 39 | 54.50408 | 60.294 |
| 441 | 1 | 13 | Long-term dynamics of bird populations in birch forests of Ilmen Nature Reserve during the breeding period individuals / km2 | Birds | Forest | 39 | 54.50408 | 60.294 |
| 469 | 16 | 20.38 | A multi-decade time series of kelp forest community structure at the California Channel Islands. | Benthos | Kelp forest | 40 | 33.851 | -119.6 |
| 473 | 26 | 12.58 | Fourteen years of mapped permanent quadrats in a northern mixed prairie | grasses | Northern mixed prairie | 41 | 46.317 | -105.8 |
| 475 | 1 | 13 | Structure and dynamics of a passerine bird community in a spruce-dominated boreal forest | Birds | Boreal forest | 42 | 63.417 | 10.5 |
| 478 | 1 | 37 | Long term study of the stream ecosystems in the Breitenbach | Insects | Central European streams | 43 | 50.67 | 9.6289 |
| 483 | 5 | 24 | ITEX Dataset 5 - Teberda (Malaya Alpine-Snowbed and Geranium Hedysarum Meadow) | plants | Alpine | 44 | 43.27 | 41.41 |
| 497 | 19 | 25.89 | ITEX Dataset 19 - Teberda (Festuca Varia Grassland, Malaya Alpine Lichen-Heath) | plants | Alpine | 44 | 43.27 | 41.42 |
| 499 | 1 | 12 | Macrobenthos monitoring at long-term monitoring stations in the Belgian part of the North Sea between 1979 and 1999 | Macrobenthos | Coastal habitats | 45 | 51.485 | 2.8953 |

