## Supplementary material for "Deviations from dynamic equilibrium in ecological communities worldwide": online appendix: add_bbs_data.m

```
load('BBS_2015_2.mat')

%% preliminary reshaping:

data_bbs_tmp=struct;
for si=1:length(sites)
    
    ind=si; %site index in the 'sites' database
    data_bbs_tmp(si).sites_ind=ind;
    data_bbs_tmp(si).route_code=sites(ind).route_code;
    data_bbs_tmp(si).years=sites(ind).years_route;

    
    %prepare dynamics data matrix:
    years_existing=ismember(sites(ind).years_route(1):sites(ind).years_route(end),sites(ind).years_route)'; %out of the entire range
    dyn=nan(length(years_existing),length(sites(ind).tot_abundance(1,:)));
    species_existing=sites(ind).tot_abundance(1,:);
    for sp=1:length(species_existing)
        dyn(years_existing,sp)=sites(ind).species(species_existing(sp)).ts(2,:)';
    end
    dyn(dyn>0)=1;
    data_bbs_tmp(si).dyn=dyn;
    
    data_bbs_tmp(si).observer=nan(length(years_existing),1);%
    data_bbs_tmp(si).observer(years_existing)=sites(ind).observer;
        
    data_bbs_tmp(si).sps=species_existing';

    data_bbs_tmp(si).sr = sum(data_bbs_tmp(si).dyn,2);
end

%% now add to a 'data-like' data structure:

%the output:
data_from_bbs_counter = 1;
data_from_bbs = struct;

for ss=1:length(data_bbs_tmp)
    ['prep2 ' num2str(ss)]
    
    len_max = length(data_bbs_tmp(ss).sr);
    if len_max >=thresh_years %otherwise there's nothing that can be done!
                
        %look for the start and end coordinates
        for ll=len_max:-1:thresh_years %run over different lengths of sequence, beginning with longest

            %run over all possible sequences of this length:
            missing_years=nan(1,len_max); %how many missing years in this sequence (only record acceprable sequences, without too many missing years)
            seq_years_start=nan(1,len_max); %record the start and end years of each sequence
            seq_counter=1;
            
            for y1=1:(len_max-ll+1) %run over possible starting years
                seq_years = y1:(y1+ll-1); %the sequence of years to be considered
                
                %check four conditions:
                %1. number of species in this period >= thresh_species
                %2. number of missing years <= length*thresh_missing
                %3+4. first and last years are not NA.
                
                if (sum(nansum(data_bbs_tmp(ss).dyn(seq_years,:))>0) >= thresh_species) && (sum(isnan(data_bbs_tmp(ss).sr(seq_years))) < ll*thresh_missing) && (sum(isnan(data_bbs_tmp(ss).dyn(seq_years(1),:))) == 0) && (sum(isnan(data_bbs_tmp(ss).dyn(seq_years(end),:))) == 0)

                    %now that this works, check that the changes in
                    %composition and richness are not too big:
                    
                    take_inds = (~isnan(data_bbs_tmp(ss).sr)) & (ismember(1:len_max,seq_years))' ; %subset of non-NAN years in range of 'seq_years'
                    obs_change_index=(diff(data_bbs_tmp(ss).observer(take_inds)))~=0;

                    if sum(obs_change_index) > 0
                        dyn_reduced = data_bbs_tmp(ss).dyn(take_inds,:);
                        sr_changes=abs(diff(data_bbs_tmp(ss).sr(take_inds)));
                        [~,p_richness]=ttest2(sr_changes(obs_change_index),sr_changes(~obs_change_index),'Tail','right');

                        %Jaccard:
                        j_changes=zeros(1,(length(dyn_reduced(:,1))-1));
                        for t=1:(length(j_changes))
                            j_changes(t)=sum(abs(dyn_reduced(t+1,:)-dyn_reduced(t,:)))/(sum(abs(dyn_reduced(t+1,:)-dyn_reduced(t,:)))+sum((dyn_reduced(t+1,:)+dyn_reduced(t,:))==2));
                        end
                        [~,p_composition]=ttest2(j_changes(obs_change_index),j_changes(~obs_change_index),'Tail','right');

                        if (p_richness > 0.05) && (p_composition > 0.05)
                            take_this = 1;
                        else
                            take_this = 0;
                        end
                    else
                        take_this = 1;
                    end
                    
                    %now, if this sequence is OK, add it to the list of
                    %sequences by recording its start and also the number
                    %of missing years in it:
                    
                    if take_this == 1
                        missing_years(seq_counter) = sum(isnan(data_bbs_tmp(ss).sr(seq_years)));
                        seq_years_start(seq_counter) = y1;
                        seq_counter = seq_counter + 1;
                    end
                    
                end
            end
            
            if seq_counter > 1 %if some sequence of length ll was found
                f = find(missing_years == min(missing_years));
                %take the first of f:

                data_from_bbs(data_from_bbs_counter).PLOT = data_bbs_tmp(ss).route_code;
                data_from_bbs(data_from_bbs_counter).dyn = data_bbs_tmp(ss).dyn(seq_years_start(f(1)):(seq_years_start(f(1))+ll-1),:);
                %remove empty columns:
                find_empty = nansum(data_from_bbs(data_from_bbs_counter).dyn) == 0;
                data_from_bbs(data_from_bbs_counter).dyn(:,find_empty) = [];
                
                data_from_bbs(data_from_bbs_counter).sps = data_bbs_tmp(ss).sps;
                data_from_bbs(data_from_bbs_counter).sps(find_empty) = [];
                
                year_start = data_bbs_tmp(ss).years(1) + seq_years_start(f(1)) - 1;
                data_from_bbs(data_from_bbs_counter).years = year_start:(year_start + ll - 1);
                data_from_bbs(data_from_bbs_counter).years = data_from_bbs(data_from_bbs_counter).years(~isnan(data_from_bbs(data_from_bbs_counter).dyn(:,1)))';
                data_from_bbs_counter = data_from_bbs_counter + 1;
                break
            end
            
        end
        
    end
end

%% add BBS to the big data structure:

initial_ind = length(data);
for ss = 1:length(data_from_bbs)

    data(initial_ind+ss).study_id = 1000;
    data(initial_ind+ss).lat = NaN;
    data(initial_ind+ss).lon = NaN;
    data(initial_ind+ss).plot = num2str(data_from_bbs(ss).PLOT);
    data(initial_ind+ss).toofew = 0;
    data(initial_ind+ss).years = data_from_bbs(ss).years;
    data(initial_ind+ss).diff_max = 1;
    data(initial_ind+ss).rarefied = 0;
    data(initial_ind+ss).dyn = data_from_bbs(ss).dyn;
    data(initial_ind+ss).sps = data_from_bbs(ss).sps;
end
```
