## Supplementary material for "Deviations from dynamic equilibrium in ecological communities worldwide": online appendix: add_noise.m

```
%noise1  is the noise used for the study level analysis (normalized by
%sqrt). noise2 is the noise used for the analysis at the site level, which
%is normalized by nothing for all studies but Konza, where it is normalized
%by initial richness.
'computing noise and trends'

for ss = 1:length(data)
    
    if data(ss).diff_max == 1 %if there aren't rarefac_ittrlicates
             
        ts = sum(data(ss).dyn,2); %time series of richness
        tsx = (1:length(ts))'; %the 'x' axis of the time series
        
        %cut time series:
        cut_years = ceil(cut_percent*length(ts)); %number of years to cut
        tsc = ts((cut_years+1):(end-cut_years));
        tsxc= (1:length(tsc))';
        
        %compute noise:
        data(ss).noise1 = nanstd(diff(ts)./sqrt(ts(1:end-1)));
        data(ss).noisec1 = nanstd(diff(tsc)./sqrt(tsc(1:end-1)));
        
        if data(ss).study_id~=355
            data(ss).noise2 = nanstd(diff(ts));
            data(ss).noisec2 = nanstd(diff(tsc));
        else
        	data(ss).noise2 = nanstd(diff(ts)./(ts(1:end-1)));
            data(ss).noisec2 = nanstd(diff(tsc)./(tsc(1:end-1)));
        end    
        
        %compute slope:
        [sen, ~] = TheilSen([tsx ts]);
        data(ss).sen_norm = sen/nanmean(ts);
        
        [senc, ~] = TheilSen([tsxc tsc]);
        data(ss).senc_norm = senc/nanmean(tsc);
                
    else %there are several samples, data is rarefied.
        noise2s = zeros(1,rarefac_ittr);
        noise1s = zeros(1,rarefac_ittr);
        sen_norms = zeros(1,rarefac_ittr);
        
        noisec2s = zeros(1,rarefac_ittr);
        noisec1s = zeros(1,rarefac_ittr);
        senc_norms = zeros(1,rarefac_ittr);
        
        for ii=1:rarefac_ittr
            
            ts = sum(data(ss).dyn{ii},2);
            cut_years = ceil(cut_percent*length(ts));
            
            tsx = (1:length(ts))';
            tsc = ts((cut_years+1):(end-cut_years));
            tsxc= (1:length(tsc))';

            noise1s(ii) = nanstd(diff(ts)./sqrt(ts(1:end-1)));
            noisec1s(ii) = nanstd(diff(tsc)./sqrt(tsc(1:end-1)));
            
            if data(ss).study_id~=355
                noise2s(ii) = nanstd(diff(ts));
                noisec2s(ii) = nanstd(diff(tsc));
            else
                noise2s(ii)= nanstd(diff(ts)./(ts(1:end-1)));
                noisec2s(ii)= nanstd(diff(tsc)./(tsc(1:end-1)));
            end

            [sen, ~] = TheilSen([tsx ts]);
            sen_norms(ii) = sen/nanmean(ts);
            [senc, ~] = TheilSen([tsxc tsc]);
            senc_norms(ii) = senc/nanmean(tsc);
        end
        
        %average values:
        data(ss).noise2 = mean(noise2s);
        data(ss).noise1 = mean(noise1s);
        data(ss).sen_norm = mean(sen_norms);
        
        data(ss).noisec2 = mean(noisec2s);
        data(ss).noisec1 = mean(noisec1s);
        data(ss).senc_norm = mean(senc_norms);
        
    end
end

%now average over different sites:

studies = unique([data.study_id]);

for st = 1:length(studies)
    sites_inds = find(([data.study_id] == studies(st))); %indexes of sites
    
    data_s(st).noise2 = mean([data(sites_inds).noise2]);
    data_s(st).noise1 = mean([data(sites_inds).noise1]);
    data_s(st).sen_norm = mean(abs([data(sites_inds).sen_norm]));
    data_s(st).noisec2 = mean([data(sites_inds).noisec2]);
    data_s(st).noisec1 = mean([data(sites_inds).noisec1]);
    data_s(st).senc_norm = mean(abs([data(sites_inds).senc_norm]));

end
```
