## Supplementary material for "Deviations from dynamic equilibrium in ecological communities worldwide": online appendix: add_shiran.m

```
load('data_shiran.mat')

initial_ind = length(data);

for ss = 1:length(data_shiran)
    
    
    data(initial_ind+ss).study_id = 1000 + data_shiran(ss).set_num;
    data(initial_ind+ss).lat = NaN;
    data(initial_ind+ss).lon = NaN;
    data(initial_ind+ss).plot = num2str(data_shiran(ss).tab_num);
    data(initial_ind+ss).toofew = 0;
    data(initial_ind+ss).years = data_shiran(ss).years;
    data(initial_ind+ss).diff_max = 1;
    data(initial_ind+ss).rarefied = 0;
    data(initial_ind+ss).dyn = data_shiran(ss).dyn;
    
    data(initial_ind+ss).dyn(~isnan(data(initial_ind+ss).dyn)) = data(initial_ind+ss).dyn(~isnan(data(initial_ind+ss).dyn))>0;
    data(initial_ind+ss).dyn(:,nansum(data(initial_ind+ss).dyn) == 0) = [];
end
```
