## Supplementary material for "Deviations from dynamic equilibrium in ecological communities worldwide": online appendix: Analysis.m

```
%% Quality criteria:

thresh_missing=0.15; %the maximal proportion of missing years
thresh_years = 10; %the minimal number of years
thresh_species = 20; %the minimal number of species observed overall in a site
thresh_good_month = 0.9; % if a month is present at this proportion of years or above, we will throw away every year with this month missing.
month_value = 0.3; % used to compute the optimal sequence of years. 
resamps = 20000; %total number of synthetic communities to generate per site
rarefac_ittr=50; %number of rarefaction randomizations performed (when needed)
thresh_col_ex = 1; %the minimal number of species that have at least two events (colonization and then extinction or vice versa).
cut_percent = 0.06; %the proportion of the time series that needs to be cut on the sides.

%% Now, generate the 'data' data structure from the three sources:

data=struct;

%put all data into the same format:
generate_data_BioTIME;
add_bbs_data;
add_shiran;

%minor filtering:
data( ([data.diff_max] > 1) & ~([data.study_id] == 355) ) = []; %remove all rarefied sites, with the exception of sites within the Konza Prairie dataset

%generate statistics:
calculate_stats
data(isnan([data.var_col])) = []; %remove single observation with nan in varcol.
data_s(isnan([data_s.var_col])) = []; %remove case with insufficeint dynamics
data(isnan([data.var_col])) = []; %remove case with insufficeint dynamics
%analyze_noise %this is a script performing a first analysis of the scaling
%of fluctuations across sites
add_noise

%perform final analysis:
generate_data_tables;
compute_models

save('results.mat','data','data_s')

%plot:
generate_stats_hist2
plot_dsc2
```
