## Supplementary material for "Deviations from dynamic equilibrium in ecological communities worldwide": online appendix: analyze_months3.m

```
function [table_new, months_inds] = analyze_months3(table_original, thresh_years, thresh_missing, thresh_species, thresh_good_month, mv)


%thresh_years=10;
%thresh_missing=0.15; %no more then 10% missing years!
%thresh_good_month=0.9;
%mv = 0.3; %value of a month


%create a year(row, older first)-month(column) table. This table considers
%also 'nearest months' as existing if a given month was in the data.
[~,month_year] = calc_month_year3(table_original); 

months_inds = NaN; %default

% remove years that are "holes":
% i.e: a year where at least two good months are missing. 'Good months' are
% defined as month that occur in 'thresh_good_month' or above of years.

good_months = double((sum(month_year)/length(month_year(:,1))) > thresh_good_month); %months above threshold
rem_years = good_months * abs(month_year - 1).' > 1; %which years to remove?
years_all=min(table_original.YEAR):max(table_original.YEAR);
table2 = table_original(~ismember(table_original.YEAR,years_all(rem_years)),:); %new table, with removed years
[month_year_extended,month_year_limited] = calc_month_year3(table2); %two tables, one that considers also adjaent months and one that does not. 

%go over all combinations of months and decide what to take:
months_table = zeros((2^12 -1 ),4); 
%each row: longest sequence of different combinations.
%Columns:
%1. number of years
%2. number of missing years
%3. number of months included
%4. first year

%which months actually occur in the data (not nescesarily every year)?
months_data = str2double(cellstr(unique(table2.MONTH)));
months_data = months_data((months_data > 0) & (months_data < 13));

for ii=1:2^12 -1
    months_take = de2bi(ii,12); %which months should be considered?
    months_table(ii,3) = length(intersect(find(months_take),months_data)); %make sure to count months correctly, disregarding months that do not really occur in the data.
    sum_month_extended = sum((month_year_extended(:,logical(months_take))),2)';
    sum_month_limited = sum((month_year_limited(:,logical(months_take))),2)';
    seq_now  = double(sum_month_extended == sum(months_take)); %the sequence of years when the number of months equals the number of months in the time span considered (e.g., years with four months if we consider the interval Jan-Apr)
    seq_now(sum_month_limited == 0) = 0; %make sure there are actually any months in these years!
    [seq_years,months_table(ii,2)] = longest_seq4(seq_now,thresh_missing,thresh_years);
    if ~isnan(sum(seq_years))
        months_table(ii,1) = seq_years(2)-seq_years(1)+1;
        months_table(ii,4) = seq_years(1);
    end
end

%finally, find the best combination of months and years. Take the
%combination with the hights value of years*(months^mv), where mv would
%typically be~0.3. In this case, 50 years with 5 months will be better then
%40 years with 10 months but worse than 40 years with 11 months. among
%options with similar values, pick the first among those with the fewest
%missing years.

vals = months_table(:,1).*(months_table(:,3).^mv);

%are there any good sequences at all?
if sum(vals)>0
    
    best_vals = find(vals == max(vals)); %indexes of best sequences
    if length(best_vals) == 1; %only one such best sequence
        take_months = de2bi(best_vals,12);
        
        %which indexes of years to take?
        ind_first = months_table(best_vals,4);
        ind_last = ind_first + months_table(best_vals,1) - 1;
    else %choose the one with the most months.
        months_table2 = months_table(best_vals,:);

        f = find(months_table2(:,2) == min(months_table2(:,2)));
        take_months = de2bi(best_vals(f(1)),12);
        
        %which indexes of years to take?
        ind_first = months_table2(f(1),4);
        ind_last = ind_first + months_table2(f(1),1) - 1;
    end
    
    %filter years:
    seq_initial  = sum(month_year_extended(:,logical(take_months)),2)'; 
    %remove unwarranted years:
    seq_initial([1:(ind_first-1) (ind_last+1):length(seq_initial)]) = 0; %nullify years that are out of the boundaries    
    seq_initial = (seq_initial == max(seq_initial)); %take only the years with all months present
    years_all_new=min(table2.YEAR):max(table2.YEAR);
    years_take = years_all_new(seq_initial); %transform indexes into actual years
    
    %leave only filtered months:
    months_inds = intersect(find(take_months),months_data); %only those with observations!
    %table_new = table2(ismember(str2num(char(table2.MONTH)),months_inds),:);
    table_new = table2(ismember(str2double(cellstr(table2.MONTH)),months_inds) & ismember(table2.YEAR,years_take),:);
    
    if (height(table_new) == 0) || (length(unique(table_new.ID_SPECIES)) < thresh_species) %check if we did not get an empty table and if there are enoughth species
        table_new = NaN;
    end

else %no good sequences
    table_new = NaN;
end

end 
```
