## Supplementary material for "Deviations from dynamic equilibrium in ecological communities worldwide": online appendix: analyze_noise.m

```
%analyzes the scaling of year to year fluctuations with initial richness

%there are several alternative options:

%inds = find([data.study_id] ~= 1000); %all data excluding BBS
%inds = find([data.study_id] == 163); %NP Fish
%inds = find([data.study_id] == 355); %Konza Praerie 
%inds = find([data.study_id] == 428); %SKagerrak Fish 
inds = find([data.study_id] == 1000); %BBS


pre_size = 1000000; %prealocation
counter = 1;

delta_s = zeros(1,pre_size); %changes in richness, centered and squared
s0 = zeros(1,pre_size); %initial richenss


for ss = inds
    
    if data(ss).diff_max == 1    
        sr_ts = sum(data(ss).dyn,2); %richness time series
    else
        sr_ts = sum(data(ss).dyn{1},2); %richness time series
    end
        d = diff(sr_ts)';
        d = d-nanmean(d);
        diffs = d.^2; %change in richness
        srs = sr_ts(1:end-1)'; %initial richness
        take = ~isnan(diffs); %which indexes to take?
        sum_take = sum(take); %how many values to add?
        
        %add data:
        delta_s(counter:(counter+sum_take-1)) = diffs(take);
        s0(counter:(counter+sum_take-1)) = srs(take);
        counter =   counter + sum_take;

end

%remove excess data:
delta_s = delta_s(1:counter-1);
s0 = s0(1:counter-1);
     
cftool(s0,delta_s)
```
