## Supplementary material for "Deviations from dynamic equilibrium in ecological communities worldwide": online appendix: calc_month_year3.m

```
function [month_year2,month_year] = calc_month_year3(r_now)
%create a year(row, older first)-month(column) table
%this version exports both tables - the 'expanded' table (2) and the
%original one. the 'expanded' table puts '1' when the year before or after
%had any data

uy = unique(r_now.YEAR);
uy2 = uy(1):uy(end); %the full range of years, including missing_years
month_year = zeros(length(uy2),12);

for yy = 1:length(uy2)
   
   months1 = unique(r_now.MONTH(r_now.YEAR == uy2(yy))); %returns a cell array of all unique months
   months2 = str2double(cellstr(months1));% now we have an array of numbers 
   months2 = months2(months2>0 & months2<13);
   month_year(yy,months2)=1; %change the existing months to 1;
end

%the next line is a bit complicated. It adds to each month_year '1' if
%there is a '11 in the month before or prior. to enforce the cyclic
%boundary conditions, we add two matrices. The first one answers the
%question: is the month after you '1'? the second answers: 'is the month
%brfore you '1'? unknown months (before the first month in the data and
%after the last month in the data) get 0.
month_year2 = month_year + [month_year(:,2:12) [month_year(2:end,1) ; 0]] + [[0 ; month_year(1:end-1,12)] month_year(:,1:11)];
month_year2 =double(month_year2 > 0);

end
```
