## Supplementary material for "Deviations from dynamic equilibrium in ecological communities worldwide": online appendix: calculate_stats.m

```
%run over all studies and calculate the site and study level statistics and
%their significance:

%the code runs a for loop over all studies. then, within every study, a
%parallel for (parfor) over different sites. Hence, before the parfor, a
%set of temporary variables is declared. The script computes the statistics
%and p values for every site and then assignes them back to the sites.
%(this is essential because of the requitements of parfor). Finally, the
%average ststistics are calculated for every study. p values at the study
%level are computed the following way: for studies with many sites (above
%'thresh_t_test', a simple t test is used. if there aren't so many sites,
%the resampled statistics are exported by the function 'test_all4_mex' and
%are then used to test whether the average in a study differs from 0.

tic

thresh_t_test = 200;

studies = unique([data.study_id]);
data_s = struct; %study level data structure of results

resamps_per_rarefac = round(resamps/rarefac_ittr);

for st = 1:length(studies)
    
    st
    
    sites_inds = find(([data.study_id] == studies(st))); %indexes of sites
    total_sites = length(sites_inds);
    
    %the statistics for each site and their respective p_values:
    djs = nan(total_sites,1);
    var_exs = nan(total_sites,1);
    var_cols = nan(total_sites,1);
    c_means = nan(total_sites,1);
    dss = nan(total_sites,1);
    dscs = nan(total_sites,1);
    dj_pvals = nan(total_sites,1);
    var_ex_pvals = nan(total_sites,1);
    var_col_pvals = nan(total_sites,1);
    c_mean_pvals = nan(total_sites,1);
    ds_pvals = nan(total_sites,1);
    dsc_pvals = nan(total_sites,1);
    toofews = zeros(total_sites,1);
    avg_srs = nan(total_sites,1);
    
    if total_sites < thresh_t_test %if there aren't too many sites
        export_res = true;
    else
        export_res = false;
    end
    
    if export_res == true %if there aren't too many sites, export resampled data to compute site level p values
        dj_res = nan(total_sites,resamps);
        var_ex_res = nan(total_sites,resamps);
        var_col_res = nan(total_sites,resamps);
        c_mean_res = nan(total_sites,resamps);
        ds_res = nan(total_sites,resamps);
        dsc_res = nan(total_sites,resamps);
    end
    
    parfor si = 1:length(sites_inds)
        %check that the data has at least one prence-absence-presence time
        %series and one absence-presence-absence time series, or stop analysis of this site:
        this_site_ind = sites_inds(si);
        
        if data(this_site_ind).diff_max == 1 %no rarefaction
            dyn_rem_nan = data(this_site_ind).dyn(~isnan(data(this_site_ind).dyn(:,1)),:); %remove nans from 'dyn' to compute the number of changes
            dd=diff(dyn_rem_nan);
            sdd_abs=nansum(abs(dd)); %numer of events in a time series.
            %require at least 'thresh_col_ex' coloniaztion events or
            %extinction events in time series with at least 3 sections.
            if sum((sdd_abs>=2)) < thresh_col_ex
                toofews(si).toofew = 1; %flag that there aren't enougth dynamics to perform the analysis.
                continue
            else
                if export_res == true
                    [c_means(si),c_mean_pvals(si),var_exs(si),var_ex_pvals(si),var_cols(si),var_col_pvals(si),dss(si),ds_pvals(si),dscs(si),dsc_pvals(si),djs(si),dj_pvals(si),c_mean_res(si,:),var_ex_res(si,:),var_col_res(si,:),ds_res(si,:),dsc_res(si,:),dj_res(si,:)] = test_all4_mex(data(this_site_ind).dyn,resamps,cut_percent,true);
                else
                    [c_means(si),c_mean_pvals(si),var_exs(si),var_ex_pvals(si),var_cols(si),var_col_pvals(si),dss(si),ds_pvals(si),dscs(si),dsc_pvals(si),djs(si),dj_pvals(si),~,~,~,~,~,~] = test_all4_mex(data(this_site_ind).dyn,resamps,cut_percent,false);
                end
                
                avg_srs(si) = nanmean(sum(data(this_site_ind).dyn,2));
            end 
        else %need to rarify:
            
            %prepare input:
            if export_res == true
                c_mean_res_add = nan(rarefac_ittr,resamps_per_rarefac);
                var_ex_res_add = nan(rarefac_ittr,resamps_per_rarefac);
                var_col_res_add = nan(rarefac_ittr,resamps_per_rarefac);
                ds_res_add = nan(rarefac_ittr,resamps_per_rarefac);
                dsc_res_add = nan(rarefac_ittr,resamps_per_rarefac);
                dj_res_add = nan(rarefac_ittr,resamps_per_rarefac);
            end
        
            c_means_add = nan(rarefac_ittr,1);
            c_mean_pvals_add = nan(rarefac_ittr,1);
            var_exs_add = nan(rarefac_ittr,1);
            var_ex_pvals_add = nan(rarefac_ittr,1);
            var_cols_add = nan(rarefac_ittr,1);
            var_col_pvals_add = nan(rarefac_ittr,1);
            dss_add = nan(rarefac_ittr,1);
            ds_pvals_add = nan(rarefac_ittr,1);
            dscs_add = nan(rarefac_ittr,1);
            dsc_pvals_add = nan(rarefac_ittr,1);
            djs_add = nan(rarefac_ittr,1);
            dj_pvals_add = nan(rarefac_ittr,1);
            toofews_add = zeros(rarefac_ittr,1);
            avg_srs_add = nan(rarefac_ittr,1);
            
            for rr = 1:rarefac_ittr
                dyn1 = data(this_site_ind).dyn{rr};
                
                dyn_rem_nan = dyn1(~isnan(dyn1(:,1)),:);
                dd=diff(dyn_rem_nan);
                sdd_abs=nansum(abs(dd)); %numer of events in a time series 
                %require at least 'thresh_col_ex' coloniaztion events or
                %extinction events in time series with at least 3 sections.

                if sum((sdd_abs>=2)) < thresh_col_ex
                    toofews_add(rr) = 1;
                    continue
                else
                    if export_res == true
                        [c_means_add(rr),c_mean_pvals_add(rr),var_exs_add(rr),var_ex_pvals_add(rr),var_cols_add(rr),var_col_pvals_add(rr),dss_add(rr),ds_pvals_add(rr),dscs_add(rr),dsc_pvals_add(rr),djs_add(rr),dj_pvals_add(rr),c_mean_res_add(rr,:),var_ex_res_add(rr,:),var_col_res_add(rr,:),ds_res_add(rr,:),dsc_res_add(rr,:),dj_res_add(rr,:)] = test_all4_mex(dyn1,resamps_per_rarefac,cut_percent,true);
                    else
                        [c_means_add(rr),c_mean_pvals_add(rr),var_exs_add(rr),var_ex_pvals_add(rr),var_cols_add(rr),var_col_pvals_add(rr),dss_add(rr),ds_pvals_add(rr),dscs_add(rr),dsc_pvals_add(rr),djs_add(rr),dj_pvals_add(rr),~,~,~,~,~,~] = test_all4_mex(dyn1,resamps_per_rarefac,cut_percent,false);
                    end
                    
                    avg_srs_add(rr) = nanmean(sum(dyn1,2)); %average richness
                end
            end
            
            %assign:
            c_means(si) = nanmean(c_means_add);
            c_mean_pvals(si) = nanmean(c_mean_pvals_add);
            var_exs(si) = nanmean(var_exs_add);
            var_ex_pvals(si) = nanmean(var_ex_pvals_add);
            var_cols(si) = nanmean(var_cols_add);
            var_col_pvals(si) = nanmean(var_col_pvals_add);
            dss(si) = nanmean(dss_add);
            ds_pvals(si) = nanmean(ds_pvals_add);
            dscs(si) =nanmean(dscs_add);
            dsc_pvals(si) = nanmean(dsc_pvals_add);
            djs(si) = nanmean(djs_add);
            dj_pvals(si) = nanmean(dj_pvals_add);
            toofews(si) = nanmean(toofews_add);
            avg_srs(si) = nanmean(avg_srs_add);
            
            if export_res == true
                c_mean_res(si,:) = reshape(c_mean_res_add,1,resamps);
                var_col_res(si,:) = reshape(var_col_res_add,1,resamps);
                var_ex_res(si,:) = reshape(var_ex_res_add,1,resamps);
                ds_res(si,:) = reshape(ds_res_add,1,resamps);
                dsc_res(si,:) = reshape(dsc_res_add,1,resamps);
                dj_res(si,:) = reshape(dj_res_add,1,resamps);
            end
        end
    end
    
    %after parfor, assign to data:
    
    for si = 1:length(sites_inds)
        this_site_ind = sites_inds(si);
        
        data(this_site_ind).c_mean = c_means(si);
        data(this_site_ind).c_mean_pval = c_mean_pvals(si);
        data(this_site_ind).var_ex = var_exs(si);
        data(this_site_ind).var_ex_pval = var_ex_pvals(si);
        data(this_site_ind).var_col = var_cols(si);
        data(this_site_ind).var_col_pval = var_col_pvals(si);
        data(this_site_ind).ds = dss(si);
        data(this_site_ind).ds_pval = ds_pvals(si);
        data(this_site_ind).dsc = dscs(si);
        data(this_site_ind).dsc_pval = dsc_pvals(si);
        data(this_site_ind).dj = djs(si);
        data(this_site_ind).dj_pval = dj_pvals(si);
        data(this_site_ind).toofew = toofews(si);
        data(this_site_ind).avg_sr = avg_srs(si);
        data(this_site_ind).time_span = data(this_site_ind).years(end) - data(this_site_ind).years(1) + 1;
        
        %add some further computations:
        data(this_site_ind).ds_rel = data(this_site_ind).ds/data(this_site_ind).avg_sr;
        data(this_site_ind).dsc_rel = data(this_site_ind).dsc/data(this_site_ind).avg_sr;
    end
    
    %finally, generate STUDY LEVEL summary:
    
    data_s(st).study_id = studies(st);
    data_s(st).plot_num = total_sites;
    data_s(st).avg_years = mean([data(sites_inds).time_span]);
    data_s(st).avg_diff_max = mean([data(sites_inds).diff_max]);
    data_s(st).avg_sr = mean([data(sites_inds).avg_sr]);
    
    %average statistics:
    data_s(st).c_mean = mean([data(sites_inds).c_mean]);
    data_s(st).var_ex = mean([data(sites_inds).var_ex]);
    data_s(st).var_col = mean([data(sites_inds).var_col]);
    data_s(st).ds = mean([data(sites_inds).ds]);
    data_s(st).dsc = mean([data(sites_inds).dsc]);
    data_s(st).ds_rel = mean([data(sites_inds).ds_rel]);
    data_s(st).dsc_rel = mean([data(sites_inds).dsc_rel]);
    data_s(st).dj = mean([data(sites_inds).dj]);
    
    
    %finally, community level p values:
    if export_res == true %compute the test using the exported resamples!
        data_s(st).c_mean_pval = dist_pval(mean(c_mean_res,1),mean(c_means));
        data_s(st).var_ex_pval = dist_pval(mean(var_ex_res,1),mean(var_exs));
        data_s(st).var_col_pval = dist_pval(mean(var_col_res,1),mean(var_cols));
        data_s(st).ds_pval = dist_pval(mean(ds_res,1),mean(dss));
        data_s(st).dsc_pval = dist_pval(mean(dsc_res,1),mean(dscs));
        data_s(st).dj_pval = dist_pval(mean(dj_res,1),mean(djs));
        
    else %if there are many communities, a simple t test will do!
        [~,data_s(st).c_mean_pval] = ttest(c_means);
        [~,data_s(st).var_ex_pval] = ttest(var_exs);
        [~,data_s(st).var_col_pval] = ttest(var_cols);
        [~,data_s(st).ds_pval] = ttest(dss);
        [~,data_s(st).dsc_pval] = ttest(dscs);
        [~,data_s(st).dj_pval] = ttest(djs);
    end
        
end

%remove case of NaNs:
data(isnan([data.var_col])) = [];
data_s(isnan([data_s.var_col])) = [];

toc
```
