## Supplementary material for "Deviations from dynamic equilibrium in ecological communities worldwide": online appendix: compute_models.m

```
all_names = {'All_studies', 'NP_Fish', 'Konza_Prairie', 'Skagerrak_Fish', 'NA_Birds'};

[results_dsc1,sig_dsc1] = generate_model_table(tbl_all,all_names,2,[4 5 6])

[results_ds1,sig_ds1] = generate_model_table(tbl_all,all_names,1,[4 5 6])


[results_dsc2,sig_dsc2] = generate_model_table(tbl_all,all_names,2,[4 5 6 7 13 14 15])

[results_ds2,sig_ds2] = generate_model_table(tbl_all,all_names,1,[4 5 6 7 13 14 15])

[results_dj1,sig_dj1] = generate_model_table(tbl_all,all_names,3,[4 5 6])

[results_dj2,sig_dj2] = generate_model_table(tbl_all,all_names,3,[4 5 6 7 13 14 15])

[results_dsc_noise2,sig_dsc_noise2] = generate_model_table(tbl_all,all_names,2,[11 12 7 19 20])

[results_ds_noise2,sig_ds_noise2] = generate_model_table(tbl_all,all_names,1,[11 12 7 19 20])

%%

%what is the proporstion of positive significant c_mean at the site level?

sum([data([data.c_mean_pval] < 0.05).c_mean] > 0)/length(data)

%for dsc:
sum([data([data.dsc_pval] < 0.05).dsc] < 0)/length(data)

%for dj:
sum([data([data.dj_pval] < 0.05).dj] > 0)/length(data)
```
