## Supplementary material for "Deviations from dynamic equilibrium in ecological communities worldwide": online appendix: create_dyn.m

```
function [dyn1,years,sps] = create_dyn(raw_now)
% GENERATE A SINGLE DATA TABLE FROM RAW DATA:
%input: raw_now, a chunk of the raw data
%output:
%dyn table, where each row is a year and each column is a species
%an array with the ids of species used
%an array of years in the data

years=unique(raw_now.YEAR);
sps=unique(raw_now.ID_SPECIES);

%generate empty data table (missing years will ba NaN, all other with zeros:
dyn1=nan(length(years(1):years(end)),length(sps)); 
for yy=years
    dyn1(yy-years(1)+1,:)=0;
end

year_inds=zeros(1,height(raw_now));
sps_inds=zeros(1,height(raw_now));

%find the index of every year in the data, 'skipping' over missing years
for yy=1:length(years)
    year_inds(raw_now.YEAR==years(yy)) = years(yy)-years(1)+1;
end

for sp=1:length(sps)
    sps_inds(raw_now.ID_SPECIES==sps(sp)) = sp;
end

dyn1(sub2ind(size(dyn1),year_inds,sps_inds)) = 1;

end
```
