## Supplementary material for "Deviations from dynamic equilibrium in ecological communities worldwide": online appendix: filter_studies1.m

```
function [take_study] = filter_studies1(raw, thresh_missing, thresh_years, thresh_species)

%gets a raw data table with some tiltering criteria and tells which studies
%to take. 'thresh_missing' is the maximal proportion of missing years,
%'thresh_years' is the minimal number of years and 'thresh_species' is the
%minimal number of species.

studies=unique(raw.STUDY_ID);
initial_studies_num=length(studies);

take_study=zeros(1,initial_studies_num);

min_years_with_data = thresh_years*(1- thresh_missing);

for ss=1:initial_studies_num
    
    this_study_binary_inds = raw.STUDY_ID == studies(ss);
    
    sp_num = length(unique(raw.ID_SPECIES(this_study_binary_inds)));
    years=(unique(raw.YEAR(this_study_binary_inds)));
    years_with_data = length(years);
    time_span=max(years)-min(years)+1;
    
    if (sp_num >= thresh_species) && (time_span >= thresh_years) && (years_with_data >= min_years_with_data)
        take_study(ss)=studies(ss);
    end
end

take_study=nonzeros(take_study);

end
```
