## Supplementary material for "Deviations from dynamic equilibrium in ecological communities worldwide": online appendix: filter_table2.m

```
function [table_new] = filter_table2(raw_now, thresh_missing, thresh_years, thresh_species)

%returns a filtered table, where the longest sequence with at least 'thresh_years' and at most
%'thresh_missing' missing years has been found. if no sequence is found of there
%aren't enouth species - returns 'NaN'.

%raw_now = raw2(raw2.STUDY_ID == 18,:);
%thresh_missing = 0.15;
%thresh_years = 10;
%thresh_species = 20;

sp_num = length(unique(raw_now.ID_SPECIES));

if sp_num >= thresh_species %if there are enought species (it there aren't, don't bother).

    years=unique(raw_now.YEAR);
    
    %check if there is a long enough time sequence:
    seq = years(1):years(end); %all the span of the years;
    seq2 = ismember(seq,years); %recode as '0' if the year is absent and '1' if it is present

    [seq_years,~] = longest_seq4(seq2,thresh_missing,thresh_years); %find initial and final index of the best sequecne

    if ~isnan(sum(seq_years)) %if any suitable sequence was found
        seq([1:(seq_years(1)-1) (seq_years(2)+1):length(seq)]) = 0; %remove years that are before of after the best sequence

        table_new = raw_now(ismember(raw_now.YEAR,nonzeros(seq)),:); %export
        
        if length(unique(table_new.ID_SPECIES)) < thresh_species % if the resulting sequence does not have enougth species.
            table_new = NaN; 
        end
        
    else %no sequence found, return 'NaN'
        table_new = NaN;
    end   
end

end
```
