## Supplementary material for "Deviations from dynamic equilibrium in ecological communities worldwide": online appendix: generate_data_BioTIME.m

```
% Import the raw data (takes long!):

tic
raw=readtable('BioTIMEQuery02_04_2018.csv');
toc


%% Filter studies:

%filter accoring to missing years, total length and number of species:
take_studies = filter_studies1(raw, thresh_missing, thresh_years, thresh_species);

%further remove all treates studies as well as the BBS:
treated=[224 235 243 248 270 275 276 311 321 338 340 358 362 382 455 456 457];
take_studies=take_studies(~ismember(take_studies,treated)); %remove treated
take_studies(take_studies == 195) = []; %remove the original version of the BBS

%generate a new dataset with filtered data:
raw2 = raw(ismember(raw.STUDY_ID,take_studies),:);

%% add the different sites to the 'data' dataset

counter = 1; %the counter of the different sites:

tic

for si=1:length(take_studies)
    si
    
    %first, find all unique sites in the data that pass the quality
    %threshold:
    
    study_raw = raw2(raw2.STUDY_ID == take_studies(si),:);
    sites = {};
    up = unique(study_raw.PLOT); %unique plot values
    counter_sites_in_study = 1;
    
    if length(up) > 1 %if there is more then one plot, then count tables as plots
        for uu=1:length(up)
            site_now = study_raw(strcmp(study_raw.PLOT,up(uu)),:);
            
            %is this table good enought?
            
            if test_table1(site_now, thresh_missing, thresh_years, thresh_species)
                sites{counter_sites_in_study} = site_now;
                counter_sites_in_study = counter_sites_in_study + 1;
            end
            
        end
    else %just one plot, filter according to lat-lon:
        
        l=[num2str(study_raw.LATITUDE) num2str(study_raw.LONGITUDE)];
        latlon=cellstr(l);
        ull=unique(latlon);
        
        for uu=1:length(ull)
            site_now = study_raw(strcmp(latlon,ull(uu)),:);
            if test_table1(site_now, thresh_missing, thresh_years, thresh_species)
                sites{counter_sites_in_study} = site_now;
                counter_sites_in_study = counter_sites_in_study + 1;
            end
        end
        
    end
    
    %after preparing the sites, run over them to generate a 'dyn' table of years (rows) on species (columns):
    
    for tt = 1:length(sites)
        ['prep1 ' num2str(si) ' ' num2str(tt)]
        
        site_now = sites{tt};

        %first, filter the data to find the longest time series that is up
        %to the quality standards:
        %is there a need to filter the dates?
        if length(unique(site_now.MONTH)) > 1
            [site_now ,~] = analyze_months3(site_now , thresh_years, thresh_missing, thresh_species, thresh_good_month, month_value); %this filters only the months needed!       
            if istable(site_now)
                data(counter).months_inds = unique(site_now.MONTH); %this records the months that were used
            end
        else
            site_now  = filter_table2(site_now , thresh_missing, thresh_years, thresh_species);
        end

        %are we left with any data table? if not, continue to next itteration. 
        if ~istable(site_now )
            continue
        end
        
        data(counter).study_id = take_studies(si);
        data(counter).lat = unique(site_now.LATITUDE);
        data(counter).lon = unique(site_now.LONGITUDE);
        data(counter).plot = unique(site_now.PLOT);
        %data(counter).toofew = 0; %assume data is OK, later this may be changed. This is a flag if the dataset/rarefied dataset did not have at least one colonization and extinctions

        data(counter).years = unique(site_now.YEAR);

        %is there a need to rarefy? count number of samples per year:
        samps_per_year = zeros(1,length(data(counter).years));
        for yy=1:length(data(counter).years)
            samps_per_year(yy) = length(unique(site_now.SAMPLE_DESC(site_now.YEAR == data(counter).years(yy))));
        end

        data(counter).diff_max = max(samps_per_year)/min(samps_per_year);

        %perform rarefaction?
        if data(counter).diff_max == 1 %no need!
            data(counter).rarefied = 0;

            %generate the data:
            [data(counter).dyn,~,data(counter).sps] = create_dyn(site_now);
            counter = counter + 1; %advance counter
        
        else %do perform rarefaction
            data(counter).rarefied = 1;
            dyns = cell(1,rarefac_ittr);

            parfor rr = 1:rarefac_ittr
                raw_rarefied = rarefy_raw(site_now);
                [dyns{rr},~,~] = create_dyn(raw_rarefied);
            end

            data(counter).dyn = dyns;
            counter = counter + 1; %advance counter

        end
    end
end

toc
```
