## Supplementary material for "Deviations from dynamic equilibrium in ecological communities worldwide": online appendix: generate_data_tables.m

```
%here we transform the data into data structures that can be used for
%statistical models and for figures. We generate a set of tables, the first
%one for the study level analysis, the second (with a for loop) for each of
%the four studies.

%in the first part of the table, X21, the data is standardizes, unlike the
%second part.
X21 = [[data_s.ds_rel]' [data_s.dsc_rel]' [data_s.dj]', [data_s.c_mean]', [data_s.var_ex]', [data_s.var_col]', [data_s.avg_years]', [data_s.avg_sr]' [data_s.noise1]' abs([data_s.sen_norm])' [data_s.noisec1]' abs([data_s.senc_norm])'];
X21_names = {'ds_rel', 'dsc_rel', 'dj', 'c_mean', 'var_ex', 'var_col', 'num_years','sr', 'noise', 'trend', 'noisec', 'trendc'};
X21 = zscore(X21);

X22 = [X21(:,4).*X21(:,5) X21(:,4).*X21(:,6) X21(:,4).*X21(:,7) [data_s.avg_years]' X21(:,7).*X21(:,9) X21(:,7).*X21(:,10) X21(:,7).*X21(:,11) X21(:,7).*X21(:,12)];
X22_names = {'c_mean_var_ex', 'c_mean_var_col', 'c_mean_num_years', 'ny', 'years_noise', 'years_trend', 'years_noisec', 'years_trendc'};

X2 = [X21 X22];
tbl1 = array2table(X2, 'VariableNames', [X21_names X22_names]);

%now generate the tables for the studies:
tbl_all = cell(1,5);
tbl_all{1} = tbl1;

studies_large = [163 355 428 1000];

for ss=2:5
    inds = [data.study_id] == studies_large(ss-1);
    
    X1 = [[data(inds).ds_rel]' [data(inds).dsc_rel]' [data(inds).dj]' [data(inds).c_mean]' [data(inds).var_ex]' [data(inds).var_col]' [data(inds).time_span]' [data(inds).avg_sr]' [data(inds).noise2]' abs([data(inds).sen_norm])' [data(inds).noisec2]' abs([data(inds).senc_norm])'];
    X1_names ={'ds_rel', 'dsc_rel', 'dj', 'c_mean', 'var_ex', 'var_col', 'num_years', 'sr', 'noise', 'trend', 'noisec', 'trendc'};
    X1 = zscore(X1);
    
    X2 = [X1(:,4).*X1(:,5) X1(:,4).*X1(:,6)  X1(:,4).*X1(:,7)  [data(inds).time_span]' X1(:,7).*X1(:,9) X1(:,7).*X1(:,10) X1(:,7).*X1(:,11) X1(:,7).*X1(:,12)];
    X2_names = {'c_mean_var_ex', 'c_mean_var_col', 'c_mean_num_years', 'ny', 'years_noise', 'years_trend', 'years_noisec', 'years_trendc'};
        
    X = [X1 X2];
    
    tbl_all{ss} = array2table(X, 'VariableNames', [X1_names X2_names]);
    
end
```
