## Supplementary material for "Deviations from dynamic equilibrium in ecological communities worldwide": online appendix: generate_hist_now4.m

```
function [] = generate_hist_now4(effect_size, sig, position,tpx,col_sig,col_nonsig)

%tpy is text position on y axis

[c,edges] = histcounts(effect_size);
max_count = max(c);
c = c/max_count;
x = edges(1)+(edges(2)-edges(1))*(0.499:1:length(edges(2:end)));
hold on
b = bar(x,c+ position,'hist');
b.FaceColor = col_nonsig;
b.LineStyle = 'none';
b.Vertices(b.Vertices==0) = position;


c2 = histcounts(effect_size(sig < 0.05),edges);
c2 = c2/max_count;
b2 = bar(x,c2  + position,'hist');
b2.Vertices(b2.Vertices==0) = position;
b2.FaceColor = col_sig;
b2.LineStyle = 'none';

text(tpx,position+0.5,[num2str(100 * round(sum(sig < 0.05)/length(sig),3)) '%'],'FontName','Times New Roman','FontSize',18, 'Color',col_sig)


end
```
