## Supplementary material for "Deviations from dynamic equilibrium in ecological communities worldwide": online appendix: generate_model_table.m

```
function [stats_table,sig_table] = generate_model_table(input_table,studies_names,rv,pv)

%this script/function generates two tables that summarizes the effect sizes
%and significance of an input tables of variables. rv is the response
%variable, pv is the vector of explanatory variables, as in the example
%below.
%input_table = tbl_all;
%studies_names = all_names;
%rv = 2;
%pv = [4 5 6];

lm_studies = cell(1,length(studies_names));

%fit a model for each s
for ss=1:length(studies_names)
    lm_studies{ss} = fitlm(input_table{ss},'ResponseVar',rv,'PredictorVars',pv,'Intercept',false); %only direct predictors
end

%ingredients of the output table
betas = zeros(length(pv),length(studies_names));
ps = zeros(length(pv),length(studies_names));
more_stats = zeros(2,length(studies_names));

for ss=1:length(studies_names)
    betas(:,ss) = lm_studies{ss}.Coefficients.Estimate;
    ps(:,ss) = lm_studies{ss}.Coefficients.pValue;
    more_stats(1,ss) = lm_studies{ss}.DFE;
    more_stats(2,ss) = lm_studies{ss}.Rsquared.Ordinary;
end

more_stats = array2table(more_stats,'RowNames',{'DFE','R2'},'VariableNames',studies_names);
stats_table = array2table(betas,'VariableNames',studies_names,'RowNames',lm_studies{ss}.PredictorNames);
stats_table = [stats_table ; more_stats];
sig_table = array2table(ps,'VariableNames',studies_names,'RowNames',lm_studies{ss}.PredictorNames);

end
```
