## Supplementary material for "Deviations from dynamic equilibrium in ecological communities worldwide": online appendix: generate_stats_hist2.m

```
studies_large = [163 355 428 1000];
study_titles = {'N.P. Fish', 'Konza Prairie', 'Skagerrak Fish', 'N.A. Birds'};

col_nonsig_com = [0.803921580314636 0.878431379795074 0.968627452850342];
col_sig_com = [0 0 1];
col_nonsig_dataset = [0.992156863212585 0.85 0.5];
col_sig_dataset = [0.979411768913269 0.35 0];

f1 = figure();

%dsrel:
subplot1 = subplot(2,3,1,'Parent',f1);
hold(subplot1,'on');
xlim([-0.5 3])
ylim([0 8])

generate_hist_now4([data_s.dsc_rel], [data_s.dsc_pval], 6.5, 1.9, col_sig_dataset, col_nonsig_dataset)

for ss=1:4
    inds = [data.study_id] == studies_large(ss);
    generate_hist_now4([data(inds).dsc_rel], [data(inds).dsc_pval], 0.5+(ss - 1)*1.5,1.9, col_sig_com, col_nonsig_com)
end
line([0 0], [0 8],'LineWidth',2,'LineStyle','--','Color',[1 0 0])
set(subplot1,'CLim',[1 2],'FontName','Times New Roman','FontSize',18,'YTick',zeros(1,0),'LineWidth',1.5, 'Ycolor',[1 1 1]);
title('Excess richness changes (ds_r_e_l)')
text(-0.4,7.5,'a.','FontName','Times New Roman','FontSize',18,'FontWeight','bold')

%dj:
subplot1 = subplot(2,3,2,'Parent',f1);
hold(subplot1,'on');
xlim([-0.1 0.4])
ylim([0 8])

generate_hist_now4([data_s.dj], [data_s.dj_pval], 6.5,0.25, col_sig_dataset, col_nonsig_dataset)

for ss=1:4
    inds = [data.study_id] == studies_large(ss);
    generate_hist_now4([data(inds).dj], [data(inds).dj_pval], 0.5+(ss - 1)*1.5,0.25, col_sig_com, col_nonsig_com)
end
line([0 0], [0 8],'LineWidth',2,'LineStyle','--','Color',[1 0 0])
set(subplot1,'CLim',[1 2],'FontName','Times New Roman','FontSize',18,'YTick',zeros(1,0),'LineWidth',1.5, 'Ycolor',[1 1 1]);
title('Excess compositional changes (dj)')
text(-0.08,7.5,'b.','FontName','Times New Roman','FontSize',18,'FontWeight','bold')

%var ex:

subplot1 = subplot(2,3,4,'Parent',f1);
hold(subplot1,'on');
xlim([-1 3.5])
ylim([0 8])

generate_hist_now4([data_s.var_ex], [data_s.var_ex_pval], 6.5,2.1, col_sig_dataset, col_nonsig_dataset)

for ss=1:4
    inds = [data.study_id] == studies_large(ss);
    generate_hist_now4([data(inds).var_ex], [data(inds).var_ex_pval], 0.5+(ss - 1)*1.5,2.1 , col_sig_com, col_nonsig_com)
end
line([0 0], [0 8],'LineWidth',2,'LineStyle','--','Color',[1 0 0])
set(subplot1,'CLim',[1 2],'FontName','Times New Roman','FontSize',18,'YTick',zeros(1,0),'LineWidth',1.5, 'Ycolor',[1 1 1]);
title('Extinction uniformity (var_e_x)')
text(-0.8,7.5,'c.','FontName','Times New Roman','FontSize',18,'FontWeight','bold')


%var col:

subplot1 = subplot(2,3,5,'Parent',f1);
hold(subplot1,'on');
xlim([-1 3.5])
ylim([0 8])

generate_hist_now4([data_s.var_col], [data_s.var_col_pval], 6.5,2.3, col_sig_dataset, col_nonsig_dataset)

for ss=1:4
    inds = [data.study_id] == studies_large(ss);
    generate_hist_now4([data(inds).var_col], [data(inds).var_col_pval], 0.5+(ss - 1)*1.5,2.3, col_sig_com, col_nonsig_com)
end
line([0 0], [0 8],'LineWidth',2,'LineStyle','--','Color',[1 0 0])
set(subplot1,'CLim',[1 2],'FontName','Times New Roman','FontSize',18,'YTick',zeros(1,0),'LineWidth',1.5, 'Ycolor',[1 1 1]);
title('Colonization uniformity (var_c_o_l)')
text(-0.8,7.5,'d.','FontName','Times New Roman','FontSize',18,'FontWeight','bold')

%average cov.:

subplot1 = subplot(2,3,6,'Parent',f1);
hold(subplot1,'on');
xlim([-0.02 0.06])
ylim([0 8])

generate_hist_now4([data_s.c_mean], [data_s.c_mean_pval], 6.5,0.04, col_sig_dataset, col_nonsig_dataset)

for ss=1:4
    inds = [data.study_id] == studies_large(ss);
    generate_hist_now4([data(inds).c_mean], [data(inds).c_mean_pval], 0.5+(ss - 1)*1.5,0.04, col_sig_com, col_nonsig_com)
end
line([0 0], [0 8],'LineWidth',2,'LineStyle','--','Color',[1 0 0])
set(subplot1,'CLim',[1 2],'FontName','Times New Roman','FontSize',18,'YTick',zeros(1,0),'LineWidth',1.5, 'Ycolor',[1 1 1]);
title('Average covariance')
text(-0.015,7.5,'e.','FontName','Times New Roman','FontSize',18,'FontWeight','bold')
```
