## Supplementary material for "Deviations from dynamic equilibrium in ecological communities worldwide": online appendix: longest_seq4.m

```
function [seq_years,missing] = longest_seq4(seq_input,max_percent_missing,min_length)

% input: sequence of 0 and ones in a row vector, e.g.: 
% seq_input=[0 0 1 1 1 0 1 1 1 0 0 0 1 1 1 0 0 1 1 1 1 0 1 1];
% output: the start and end year of the following sequence: the first of
% the longest sequences where both first and last years are '1' and where
% at most 'max_percent_missing' of the years are missing. If no sequences
% at all or longer then 'min_length' are found, returns NaN.

seq_years=NaN; %default output
missing = NaN;

%find indexes of first and last years with data:
f=find(seq_input);
if ~isempty(f)
    len_max = f(end) - f(1) + 1; %maximal length of a sequence
    if len_max > min_length %if there are enought years overall    
        %look for the start and end coordinates
        for ll=len_max:-1:min_length %run over different lengths of sequence, beginning with longest

            %run over all possible sequences of this length:

            missing_years=zeros(1,len_max); %how many missing years in this sequence (only record acceprable sequences, without too many missing years)
            seq_years_start=zeros(1,len_max); %record the start and end years of each sequence
            counter=1;

            for y1=f(1):(f(end)-ll+1)

                seq_now=seq_input(y1:(y1+ll-1));
                if (sum(seq_now == 0) <= max_percent_missing*ll) && (seq_now(1) == 1)&& (seq_now(end) == 1) %if first and last years have predence and not too many missing years, take this sequence!
                    missing_years(counter) = sum(seq_now == 0);
                    seq_years_start(counter) = y1;
                    counter = counter + 1;
                end
            end

            %if sequences were found, export the first of the sequences that
            %have the least missing years

            if counter > 1
                missing_years = missing_years(1:counter-1); %shorten
                missing = min(missing_years);
                least_missing_years_start = find(missing_years == missing); %what are the indexes of the best sequences (having the fewest missing years)?
                best_start = seq_years_start(least_missing_years_start(1)); %take the first of them
                seq_years = [best_start (best_start + ll -1)];
                break
            end
        end
    end
end

end
```
