## Supplementary material for "Deviations from dynamic equilibrium in ecological communities worldwide": online appendix: null2_1ts3.m

```
function [dyn_res2]=null2_1ts3(dyn)
%generates a resample of a single species tims series. dyn is a row array
%of 0s and 1s. NaNs are kept in the same location.

dyn_res2=dyn(1,:);
d=diff(dyn(~isnan(dyn)));
seg_num=sum(abs(d))+1;

if seg_num>2 %otherwise no need to do anything, export the original

    %locate NaNs and remove them:
    dyn_original=dyn;
    nans_inds= isnan(dyn);
    nonnan_inds= ~isnan(dyn);
    dyn=dyn(~isnan(dyn));
    ld=length(dyn);
    %identify segments:

    d=diff(dyn);
    seg_num=sum(abs(d))+1;
    segments=mod(dyn(1):(dyn(1)+seg_num-1),2);
    ps=sum(segments==1);
    as=sum(segments==0);
    p_tot=sum(dyn==1);
    a_tot=sum(dyn==0);

    %resample presences:
    if ps>1
        tmp=randperm(p_tot-1);
        stops=sort(tmp(1:ps-1));
        ps_res=[stops(1) diff([stops p_tot])];
    else
        ps_res=p_tot;
    end

    %resample absences:
    if as>1
        tmp=randperm(a_tot-1);
        moves=sort(tmp(1:as-1));
        as_res=[moves(1) diff([moves a_tot])];
    else
        as_res=a_tot;
    end

    %construct time series from segments:
    dyn_res=zeros(1,ld);
    now=dyn(1);
    counter_ps=1;
    counter_as=1;
    counter_ts=1;
    for ss=1:seg_num
        if now==1 %add a segment of ones
            add=ones(1,ps_res(counter_ps));
            counter_ps=counter_ps+1;
            now=0;
        else % add asegment of zeros
            add=zeros(1,as_res(counter_as));
            counter_as=counter_as+1;
            now=1;
        end
        len=length(add);
        dyn_res(counter_ts:counter_ts+len-1)=add;
        counter_ts=counter_ts+len;
    end
    
    %now insert back the nans:
    dyn_res2=zeros(1,length(dyn_original(1,:)));
    
    dyn_res2(nonnan_inds)=dyn_res;
    dyn_res2(nans_inds)=NaN;
    
end

end
    
```
