## Supplementary material for "Deviations from dynamic equilibrium in ecological communities worldwide": online appendix: plot_dsc2.m

```
groupings = [0 14 25 1000 ; 0 100 100 1000 ; 0 20 30 1000 ; 0 30 70 1000 ; 0 15 30 1000]; %each is the lower limit that is taken in each category
study_titles_all = {'Dataset level', 'N.P. Fish', 'Konza Prairie', 'Skagerrak Fish', 'N.A. Birds'};
colors = {[0 0.4470 0.7410]  [0.8500 0.3250 0.0980]  [0.9290 0.6940 0.1250]};
xlims = {[-2 4] [-2 4] [-3 5] [-2 4] [-2 8]};
ylims = {[-2 6] [-3 4] [-3 5] [-3 4] [-4 6]};

fig2 = figure()

for ss=1:5
    subplot1 = subplot(2,3,ss,'Parent',fig2)
    hold(subplot1,'on');
    set(subplot1,'CLim',[1 2],'FontName','Times New Roman','FontSize',18,'LineWidth',1.5);
    x = tbl_all{ss}.c_mean;
    y = tbl_all{ss}.dsc_rel;
    cat = tbl_all{ss}.ny;
    
    hold on
    for cc=1:3
        take = (cat >= groupings(ss,cc)) & (cat < groupings(ss,cc+1));
        s=scatter(x(take),y(take),'filled','MarkerFaceAlpha',.2,'MarkerEdgeColor',colors{cc},'LineWidth',1);
        %alpha(s,0.5);
    end
    hold off
    title(study_titles_all(ss))
    
    if ss==1
        legend('short','intermediate','long','FontSize',14)
        ylabel('Excess richness changes (ds_r_e_l; standardized)','FontWeight','bold')
    end
    
    if ss==4
        xlabel('Average covariance (standardized)','FontWeight','bold')
    end
    
    line([-8 8],[0 0],'Color',[0.313725501298904 0.313725501298904 0.313725501298904],...
    'LineStyle','--')
    line([0 0],[-8 8],'Color',[0.313725501298904 0.313725501298904 0.313725501298904],...
    'LineStyle','--')

    xlim(xlims{ss})
    ylim(ylims{ss})
    
end
```
