## Supplementary material for "Deviations from dynamic equilibrium in ecological communities worldwide": online appendix: rarefy_raw.m

```
function [r_new] = rarefy_raw(r_now)

%generates a rarefied table.

years = unique(r_now.YEAR);
year_num = length(years);

samp_nums = zeros(1,year_num); %how many samples each year?

for yy = 1:year_num 
    samp_nums(yy) = length(unique(r_now.SAMPLE_DESC(r_now.YEAR == years(yy))));
end
min_samps = min(samp_nums); %minimal number of samples

take_rows = false(height(r_now),1);
for yy=1:year_num 
    samples_now = unique(r_now.SAMPLE_DESC(r_now.YEAR == years(yy)));
    
    %take random draw withoug replacement:
    tmp = randperm(length(samples_now));
    samples_take = samples_now(tmp(1:min_samps));
    
    take_rows((r_now.YEAR == years(yy)) & ismember(r_now.SAMPLE_DESC,samples_take)) = 1;
end

r_new = r_now(take_rows,:);

end
```
