## Supplementary material for "Deviations from dynamic equilibrium in ecological communities worldwide": online appendix: README.docx

### How to use this code

The master script that summons all other is ‘Analysis.m’. It imports and edits data from three sources:

1. BioTIME ('BioTIMEQuery02_04_2018.csv'), which is downloadable from the BioTIME website: <http://biotime.st-andrews.ac.uk/>
2. BBS, which we downloaded and assembled from the BBS site: <https://www.pwrc.usgs.gov/bbs/>
3. additional datasets we collected and appear in table S2 of the appendix.

The data is cleaned, the statistics are calculated and analyzed and then the plots are produced. Note that running this code requires 16 GB RAM and may take a few hours to complete on a good computer (as of 2019). Parallel for loops and matlab coder are used to expediate the calculations.

The generated version is then imported into PPT files for adding the pictures and frames.
