## Supplementary material for "Deviations from dynamic equilibrium in ecological communities worldwide": online appendix: test_table1.m

```
function [take_study] = test_table1(raw, thresh_missing, thresh_years, thresh_species)

%gets a raw data table of some site with some tiltering criteria and tells whether the
%table is good according to these criteria
%this is a very 'crude' filtering that does not take into account whether
%specific sequences that fit the filtering criteria actually exist.

take_study = false; %default
min_years_with_data = thresh_years*(1- thresh_missing);

sp_num = length(unique(raw.ID_SPECIES));
years=unique(raw.YEAR);
years_with_data = length(years);
time_span=max(years)-min(years)+1;

%check if there is a long enough time sequence, and has unique coordinates:

if (sp_num >= thresh_species) && (time_span >= thresh_years) && (years_with_data >= min_years_with_data) && (length(unique(raw.LATITUDE)) == 1) && (length(unique(raw.LONGITUDE)) == 1)
    take_study=true;
end

end
```
