## Supplementary material for "Deviations from dynamic equilibrium in ecological communities worldwide": online appendix: TheilSen.m

```
function [m, b] = TheilSen(data)
% Performs Theil-Sen robust linear regression on data
%
% [m b] = TheilSen(data)
%
% data: A MxD matrix with M observations. The first D-1 columns are the
%   explanatory variables and the Dth column is the response such that
%   data = [x1, x2, ..., x(D-1), y];
%
% m: Estimated slope of each explanatory variable with respect to the
%   response varuable. Therefore, m will be a vector of D-1 slopes.
% b: Estimated offsets.
%
%
% EXAMPLE:
% -------
% n=100;
% outN=round(0.2*n);
% noise = randn(n,2)*0.1; noise(randi(n*2,[outN 1]))=randn(outN,1)*5;
% data = [linspace(0,10,n)' linspace(0,5,n)'] + noise;
% Bhat = [ones(n,1) data(:,1)]\data(:,2);
% [m, b] = TheilSen(data);
% plims = [min(data(:,1)) max(data(:,1))]';
% figure
% plot(data(:,1),data(:,2),'k.',...
%     plims,plims*m+b,'-r',...
%     plims,plims*Bhat(2)+Bhat(1),'-b','linewidth',2)
% legend('Data','TheilSen','Least Squares','location','NW')
% title(sprintf('Acual Slope = %2.3g, LS est = %2.3g, TS est = %2.3g',[0.5 Bhat(2) m]))
% 
%
% Source:
%   Gilbert, Richard O. (1987), "6.5 Sen's Nonparametric Estimator of
%   Slope", Statistical Methods for Environmental Pollution Monitoring,
%   John Wiley and Sons, pp. 217�219, ISBN 978-0-471-28878-7
%
%
%
% %%% Z. Danziger October 2014 %%%
% edits Z. Danziger September 2015:
%	- updated help
%   - speed increase for 2D case
%
%

sz = size(data);
if length(sz)~=2 || sz(1)<2
    error('Expecting MxD data matrix with at least 2 observations.')
end

if sz(2)==2         % normal 2-D case
    C = nan(sz(1));
    for i=1:sz(1)
        % accumulate slopes
        C(i,i:end) = (data(i,2)-data(i:end,2))./(data(i,1) - data(i:end,1));
    end
    m = nanmedian(C(:));                        % calculate slope estimate
    
    if nargout==2
        b = nanmedian(data(:,2)-m*data(:,1));   % calculate intercept if requested
    end
    
else                % other cases
    C = nan(sz(1),sz(2)-1,sz(1));
    for i=1:sz(1)
        C(:,:,i) = bsxfun( @rdivide,data(i,end)-data(:,end),...
            bsxfun(@minus,data(i,1:end-1),data(:,1:end-1)) );       % accumulate slopes    
    end
    Cprm = reshape( permute(C,[1 3 2]), [],size(C,2),1 );           % stack layers of C to 2D
    m = nanmedian(Cprm,1);                                          % calculate slope estimate
    
    if nargout==2
        % calculate all intercepts if requested
        b = nanmedian( bsxfun(@minus,data(:,end),bsxfun(@times,m,data(:,1:end-1))) );   
    end
    
end
```
